# Phylogenomics and the origins of sharks

**DOI:** 10.64898/2026.02.13.705779

**Authors:** Chase Doran Brownstein, Thomas J. Near

## Abstract

Genomes have the capacity to drastically modify hypotheses about the relationships of species. Despite the growing availability of non-model organism genome sequences, historically contentious portions of Tree of Life remain untested using genomic data. Here, we infer the phylogeny of sharks, skates, rays, and chimaeras using the genomes of 48 species, targeting different genomic marker types. Although phylogenetic relationships of chondrichthyans are relatively consistent across analyses, different molecular markers yield conflicting results about shark monophyly. Exons support the traditional view that sharks are monophyletic, whereas ultraconserved elements and legacy nuclear markers instead suggest that the frilled and cow sharks (*Hexanchiformes*), which retain the ancestral jaw structure of cartilaginous fishes, is the sister lineage of all other sharks and rays. The resolution of sharks as monophyletic or paraphyletic has little effect on inferences of the timescale of shark evolution or the origins of key traits, such as their ancestral ecology and genome size. We tie the diversification of living cartilaginous fishes to the transformation of marine ecosystems during the middle Mesozoic Era, confirming that living shark diversity is the product of rapid ancient diversification. Consequently, our results suggest that despite uncertainty around whether sharks are monophyletic, consensus can still be reached about major evolutionary events in this iconic vertebrate lineage.

**Significance Statement:** Living sharks, skates, rays, and chimaeras form one of the three principal groups of vertebrates. These iconic animals, which include over 1200 species, are key components of marine ecosystems and have helped us reconstruct the evolution of vertebrate genomes and phenotypes. However, much work on this group has assumed that sharks are a natural group. Here, for the first time, we leverage genome-scale data to test this hypothesis. Surprisingly, we show different genome regions reject or support hypothesis that sharks form a natural group to the exclusion of skates and rays. This throws an unexpected wrench into our understanding of the relationships of some of the oldest living vertebrate clades.

## Introduction

Genome-scale data has revolutionized our understanding of the phylogeny of animals (1– 5) and their principle clades, especially vertebrates and their closest relatives among chordates (6–11, 11–15). Only recently have the historically contentious (16–21) relationships of the earliest vertebrate divergences been tested using genomic data (22, 23). This region of the vertebrate tree, which includes the successive divergences among jawed and jawless taxa and cartilaginous and bony fishes, is notable for being location of origin for numerous innovations, including the mobile jaw, paired appendages, lungs and the swim bladder, semicircular canals, and myelinated nerves (24–27). Accurate phylogenies have been crucial for resolving the stepwise evolution of these features (22, 23, 28, 29).

The over 1200 species of living cartilaginous fishes (30) last share a common ancestor that diverged from other vertebrates at least 439 million years ago (31) and include a large proportion of unique evolutionary history (32). The early fossil record of cartilaginous vertebrates (*Chondrichthyes*) provides key clues about the origins of living jawed vertebrate diversity (33–41), and the evolution of their varied reproductive modes (42) and massive genome sizes (43–47) has received much attention in the context of understanding major transitions in vertebrate genomics and life history. Yet, we still lack a comprehensive phylogeny of the *Chondrichthyes* based on genomic data. The few studies incorporating genome-wide markers to infer chondrichthyan phylogeny (43, 48, 49) do not sample all of the major orders of sharks, meaning that they do not test the essential disagreement among evolutionary trees generated using mitochondrial (50–53) and legacy nuclear (54–56) markers or morphological characters (57, 58): whether sharks and rays are reciprocally monophyletic.

Here, we leverage genome-wide marker data for 48 species representing all major living lineages of chondrichthyans to infer their phylogenetic relationships and timescale of diversification. Using multiple types of genomic markers and legacy nuclear gene data, we provide a robust resolution of chondrichthyan phylogeny with the exception of one critical node: the root of sharks, skates, and rays to the exclusion of chimaeras (*Elasmobranchii*). Phylogenies inferred from ultraconserved elements confidently reject the monophyly of sharks relative to rays and skates and show that six-gilled and frilled sharks in the clade *Hexanchiformes* are the living sister to all other elasmobranchs. However, phylogenies built using single-copy exons support the traditional hypothesis that sharks and rays are reciprocally monophyletic. By examining the proportions of individual genes and sites that support these alternative hypotheses and considering factors such as compositional bias, we demonstrate that exon sequences might be less reliable for inferring deep chondrichthyan relationships, but that the root of sharks is nonetheless a hard phylogenetic problem. Nonetheless, we show that key inferences about the evolution of sharks, including their ancestral ecology, genome size, and tempo of major lineage divergences, remain consistent even when shark paraphyly is considered. In particular, we confirm a Mesozoic diversification of chondrichthyans that included the rapid interordinal diversification of both galeomorph and squalomorph sharks during the Early and Middle Jurassic. These results illuminate the exceptionally ancient diversity contained in living sharks, rectify long-debated scenarios of jaw, genome size, life history, and ecomorphological evolution in chondrichthyans, and provide new clarity on the evolutionary history of one of three major divisions of jawed vertebrates.

## Results and Discussion

### A hard phylogenetic problem at the common ancestry of sharks and rays

Using a total of 349 ultraconserved elements (UCEs) and 840 basic universal single copy orthologs (BUSCOs) extracted from 48 chondrichthyan genomes and six osteichthyan outgroups, we conducted comprehensive phylogenomic analyses of cartilaginous vertebrates. UCEs are highly conserved regions of genomes found across taxa with ancient divergences; targeting UCEs also allows for selection of their more variable flanking regions (59). In contrast, BUSCOs are conserved protein-coding sequences (60). Consequently, targeting BUSCOs and UCEs provides us with two compositionally different sequence datasets with which to analyze and infer chondrichthyan relationships. The resulting phylogenies (Figure 1A, Figure 2; Figures S1-S12) inferred using maximum likelihood and a multispecies coalescent model are congruent across most major clades of contention in *Chondrichthyes*, supporting skates *(Rajiformes*) and torpedo rays (*Torpediniformes*) as the first two successive divergences in rays and skates (*Batoidea*), the monophyly of a clade containing sharks in the orders *Carchariniformes* and *Lamniformes*, and the monophyly of *Squalomorphii* (dogsharks, angelsharks, sawsharks, and prickly sharks) (Figure 1). Analyses of legacy molecular datasets have resolved the relationships of these lineages in many different ways (32, 43, 44, 50, 55, 56, 61–63), and so our phylogenomic analyses incorporating different genome marker datasets provide consensus regarding key chondrichthyan relationships.

**Figure 1.**
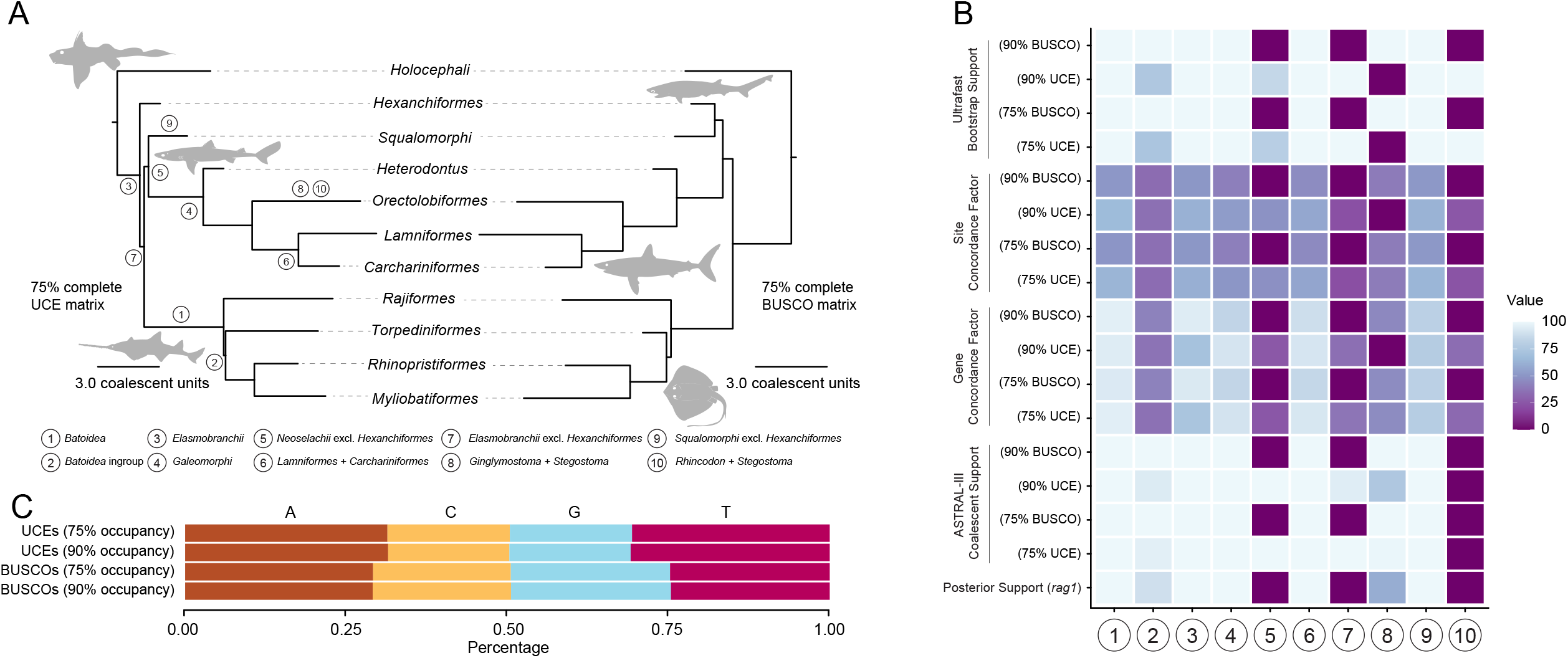
Phylogenomic analysis reveals chondrichthyan relationships. (A) Comparison of tree topologies generated using ASTRAL-III using BUSCOs and UCEs. (B) Comparison of nodal support for selected nodes across different phylogenetic methodologies and different sequence dataset completeness levels. (C) Comparisons of nucleotide base pair content across datasets.

**Figure 2.**
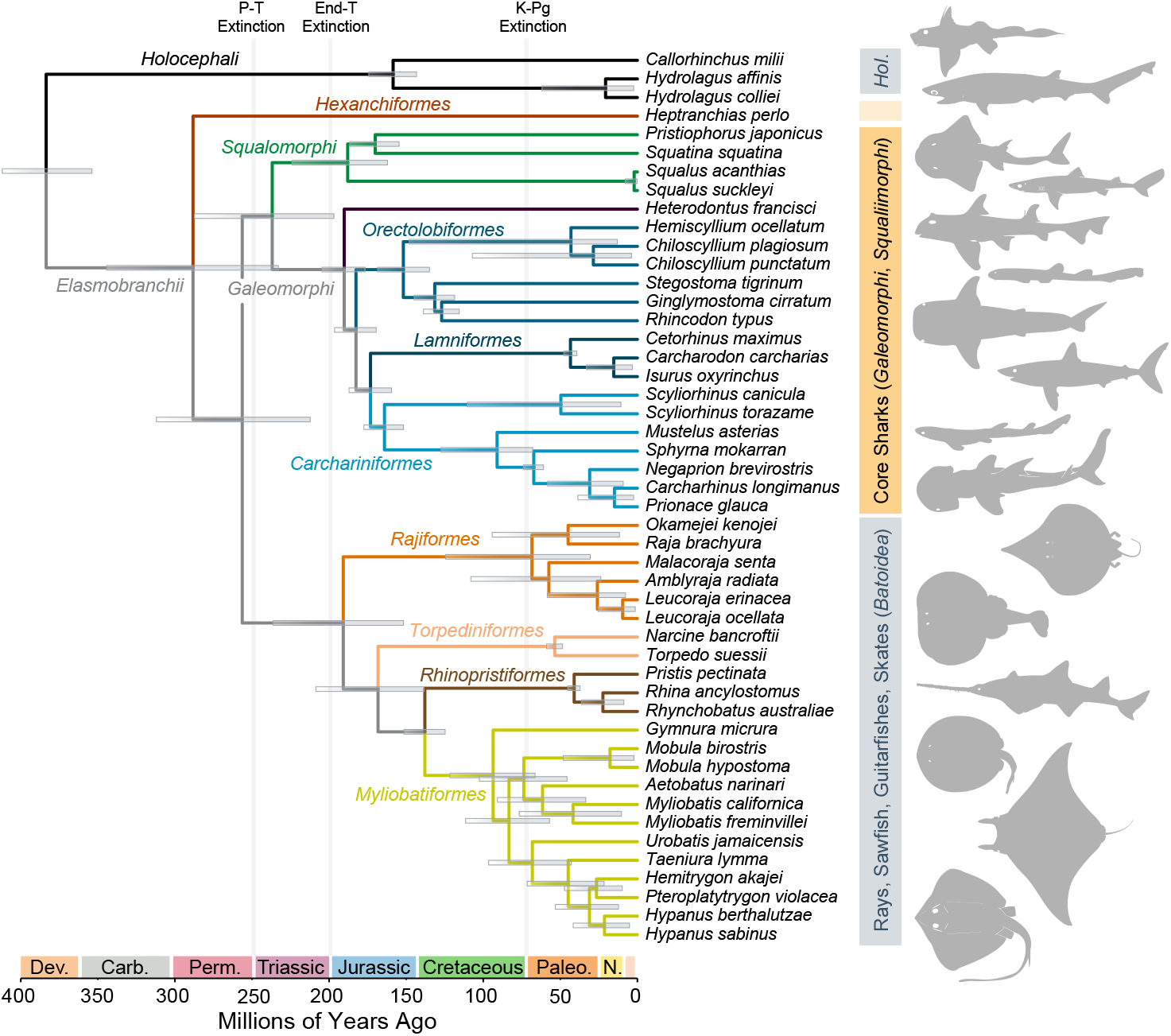
Time calibrated phylogeny of cartilaginous vertebrates. Shown is the time-calibrated phylogeny of *Chondrichthyes* made by using a Bayesian node-dating approach in BEAST 2.6.6 to calibrate the ASTRAL-III multispecies coalescent phylogeny of cartilaginous fishes based on 356 ultraconserved elements. Bars at nodes denote 95% highest posterior density intervals for divergence times. Dotted branch indicates region of ASTRAL-III phylogeny that differs from the topology generated using different partitioning approaches in IQ-TREE2. Illustrations are public domain.

Nonetheless, phylogenies inferred from UCEs and BUSCOs consistently resolve alternative topologies for the earliest divergence among sharks, skates, and rays (*Elasmobranchii*) (Figure 1). Phylogenetic analysis of UCEs results in *Hexanchiformes* (represented by *Heptranchias perlo* in our phylogeny) resolved as the sister lineage of all other sharks and rays. The *Squalomorphii*, which includes dogfishes, angelsharks, sawsharks, and pricklysharks (50, 51, 55), is resolved with weak node support as the sister to other sharks besides *Hexanchiformes* or outside a clade containing rays and skates (*Batoidea*) and all other sharks *(Galeomorphii*). In contrast, all analyses of the BUSCO dataset resolve a traditional *Squalomorphii* in which *Hexanchiformes* as the earliest diverging lineage (Figure 1A) (32, 50, 51, 56, 61–63). Interrogation of different metrics of node support across analyses shows that the resolution of *Hexanchiformes* as the sister lineage to all other sharks and rays is not the only region of phylogenetic instability; whereas monophyly of all other sharks and rays to the exclusion of *Hexanchiformes* is consistently supported by bootstrap values of 100 and high gene and site concordance factors, the monophyly of the clade consisting of all other sharks (*Galeomorphii* + *Squalomorphii*) is only moderately supported (Figure 1B). In contrast, all phylogenies inferred from BUSCOs strongly infer shark monophyly. Our alternative inferences of shark paraphyly and monophyly are robust to different levels of data completeness (Figure 1B; Figures S1-S12). Bayesian analysis of legacy nuclear sequences sampled for a larger number of chondrichthyan species (including the deeply divergent *Echinorhinus* prickly sharks) resolves sharks as paraphyletic, placing *Heptranchias perlo* as the sister taxon to other squalomorphs with weak node support (posterior = 0.51; Figure S10). Although we were unable to sample *Echinorhinus* in our phylogenomic analyses, previous phylogenies generated using mitochondrial and nuclear markers (32, 50, 55, 61) as well as genome-wide exon markers (64) consistently place *Echinorhinus* in *Squalomorphii* nested within *Squaliformes, Squatina*, and *Pristiophoriformes*, which is consistent with our phylogenetic analysis of legacy marker data (Figure S7).

Phylogenetic incongruence among different genome-wide datasets has become widely observable across major vertebrate clades such as birds (8–10, 65–72), placental mammals (7, 73–75), and teleost fishes (6, 76–79). To further identify sources of support for shark monophyly across our UCE and BUSCO datasets, we examined both the base composition of the underlying sequence data and phylogenetic support across gene trees and sites. Failure to resolve shark monophyly in molecular phylogenetic analyses of legacy nuclear markers has been attributed to GC bias (56), and phylogenetic analyses of other clades using UCEs and BUSCOs have shown that GC bias might limit the capacity of BUSCOs to infer the relationships of deeply divergent species (71). Comparisons of the nucleotide base composition of our datasets shows that the sampled BUSCOs have a considerably (3-6%) higher GC content than sampled UCEs (Figure 1C). Second, though BUSCOs have a higher gross number of parsimony informative sites (PIS) than UCEs, they also exhibit a higher degree of PIS variance per sequence alignment (Table 1). Gene and site concordance factors calculated for species trees inferred using BUSCOs also exhibit less linear relationships than those calculated for trees inferred using UCEs (Figure S13). For these reasons, we slightly favor the hypothesis supported using UCEs though acknowledge the initial divergence of elasmobranchs is a hard phylogenetic problem.

The phylogeny of sharks that we infer using UCEs would render the terms *Selachii* Gadow 1898 (80) and *Neoselachii* Compagno 1977 *(58)*, used to refer to a monophyletic shark lineage (54, 81), as junior subjective synonyms of *Elasmobranchii* Bonaparte 1838 (82). These results differ considerably from previous studies using molecular data (Figure S7), where *Hexanchiformes* is found to be the oldest-diverging clade in *Squalomorphii* and sharks are monophyletic relative to rays (50, 51, 55, 62). Our inference of paraphyly of sharks is robust to different levels of data completeness (Figure 2A, B; Figures S1-S7), and is congruent with Bayesian analysis of legacy nuclear sequences sampled for a larger number of chondrichthyan species (including the deeply divergent *Echinorhinus* prickly sharks), which fails to resolve sharks as monophyletic and places *Heptranchias perlo* as the sister taxon to other squalomorphs with weak node support (posterior = 0.51; Figure S8). Although we were unable to sample *Echinorhinus* in our phylogenomic analyses, previous phylogenies generated using mitochondrial and nuclear markers (32, 50, 55, 61) as well as genome-wide exon markers (64) consistently place *Echinorhinus* in *Squalomorphii* nested within *Squaliformes, Squatina*, and *Pristiophoriformes*, a position consistent with the phylogeny inferred from our analysis of legacy markers (Figure S7).

Shark paraphyly was previously proposed on the basis of morphological characters (83, 84), a result that was subsequently challenged several times on the basis of molecular phylogenies built using legacy nuclear and mitochondrial sequence data (50, 55, 56, 61–63). Although our analyses of BUSCOs reject the paraphyly of *Squalomorphii* relative to *Batoidea* as found in these analyses of morphological data (83, 84), the phylogenetic resolution of *Hexanchiformes* in analyses of the UCE dataset is consistent with the hypothesis that amphistylic jaw suspension is ancestral for elasmobranchs (85–87). Our results also challenge the hypothesis that the root of *Elasmobranchii* is unresolvable due to the long divergence time between elasmobranchs and holocephalans (56). A previous study of legacy nuclear markers rejected the hypothesis that *Hexanchiformes* is sister to other sharks and *Batoidea* based on the long branch lengths subtending *Elasmobranchii*, despite resolving *Hexanchiformes* in the same position as we do using UCEs (56). A similar argument was put forth with reference to the same problem in a recent preprint using genome-wide loci (88), which suggested that extensive filtering is necessary to prevent long branch attraction, heterogenous evolutionary rates, and GC content bias from inhibiting resolution of shark relationships. This hypothesis is also challenged by the resolution of older or similarly old relatiships across the animal Tree of Life using genomic data (1–3, 5), including the resolution of lungfishes and coelacanths as sarcopterygians (15, 89–95), the stepwise divergences of the major ray-finned fish lineages (6, 12, 13, 96–99) among vertebrates. In our node-dated phylogeny, we estimate that crown *Chondrichthyes* appeared 383.54 million years ago (Ma) [95% highest posterior density interval (HPD): 353.81, 412.01 Ma], *Hexanchiformes* diverged from other sharks and rays 278.09 million years ago 288.26 Ma (95% HPD: 232.86, 344.27 Ma), and all other sharks and rays share a most recent common ancestor 256.35 Ma (95% HPD: 212.24, 312.02 Ma). These results suggest that the crown age of cartilaginous vertebrates is in fact up to 40 million years younger than the age of crown-group bony vertebrates, which are known from fossils as old as the Silurian, approximately 419 million years ago (100–103).

Our hypothesis of the timescale chondrichthyan evolution based on the phylogeny inferred from UCEs supports that the interordinal divergences of elasmobranchs largely correspond to a major periods of marine faunal reorganization during the Jurassic (32, 47, 61, 63, 104, 105) (Figure 2). The exception to this is *Hexanchiformes*, which we estimate has a stem lineage that extends into the Paleozoic, highlighting this lineage as a priority for conserving ancient chondrichthyan biodiversity (32). Nearly all shark and ray interordinal divergences occur during the Early-Middle Jurassic, including a rapid diversification among shark order-level lineages (Figure 2) including *Carcharhiniformes* (hammerheads, requiem sharks, and catsharks), *Lamniformes* (white sharks, megamouth and basking sharks, and thresher sharks), and *Orectolobiformes* (whale sharks, wobbegong sharks, zebra sharks, and nurse sharks). This period coincided with Mesozoic Marine Revolution, which featured major diversifications of benthic and demersal species, as well as large predators (106–111). Phylogenomic studies of teleost fishes (6) suggest broad interordinal diversification during this time in the marine realm across depth ranges (112, 113). Similarly, transcriptomic data from coleoid cephalopods (114) indicates that the initial diversification of octopuses and squids occurred during the Triassic-Jurassic. Further, this result supports the hypothesis (61, 104) that large-bodied, pelagic sharks appeared in the Jurassic and Early Cretaceous, well after the initial diversification of most marine reptile clades in the earliest Triassic (115–118) with the notable exception of several marine crocodylomorph clades (119–123) and mosasaurs (124–126).

### The evolution of jaws in sharks and rays and the rise of pelagic planktivores

The alternative phylogenies of cartilaginous vertebrates that we present warrant a review of the evolution of key anatomical and physiological features in sharks and rays. Ancestrally, sharks and rays (*Elasmobranchii*) are inferred to have either amphistylic jaw suspension, wherein the mandibular arch is braced by the hyomandibula and there is a two-point suspension of the upper jaw-formed by the palatoquadrate-to the chondrocranium, or autodiastyly, where an additional ethmoid articulation forms a three-point suspension system (86, 87, 127, 128). Sharks in the clade *Hexanchiformes* are unique among living species in retaining amphistylic jaw suspension, and have also partially developed orbitostyly, where the palatoquadrate is also anchored to the orbital (86). These modifications to jaw suspension in sharks and rays modify the mobility of the jaws. For example, rays and skates have highly mobile euhylostylic jaws that articulate with the chondrocranium solely via the hyomandibula (87).

Hypotheses of shark phylogeny that united *Hexanchiformes* with *Squalomorphii* imply that species in the former clade represent in part a retained amphistylic condition likely found in the ancestral elasmobranch but also acquired features of orbitostylic condition found in lineages of squalomorphs (Figure 1) (83, 84, 86, 87, 128). Our phylogenomic analyses of UCEs, which infer that *Hexanchiformes* is the sister lineage of all other elasmobranchs, imply that the morphology of the jaw articulation in this clade represents the retention of the plesiomorphic condition in elasmobranchs (87). This result consequently suggests that the modified amphistyly of hexanchiform sharks represents the retention of the condition seen in many [but not all, see (38)] early-diverging elasmobranchs, such as the symmoriform pan-holocephalan (36, 129) †*Cladoselache* (87) and the pan-elasmobranch †*Phoebodus* (34).

Because we infer a Paleozoic origin for *Hexanchiformes*, it does appear that this deep-water lineage of sharks represents a survivor of ancient Paleozoic elasmobranch diversity. Previous studies that estimated the divergence time of *Hexanchiformes* from other elasmobranchs inferred a Triassic age owing to the placement of this clade within *Squalomorphii* (32, 47, 61). However, the estimated total clade age of *Hexanchiformes* in our phylogeny is younger than Devonian chondrichthyan teeth with controversial phylogenetic affinities that are somewhat similar to living hexanchiforms (130), supporting the hypothesis that these Devonian fossils represent species that converged with living hexanchiforms in dental morphology (131). Our time-calibrated phylogeny of sharks also does not exclude the possibility that the enigmatic of Permian-Jurassic †*Synechodontiformes*––variously considered a monophyletic (132, 133) or paraphyletic (81) group of early-diverging crown or stem-group sharks with dental similarities to some hexanchiforms (134)––are members of the hexanchiform total clade, on the stem of the clade containing all other elasmobranchs, or on the stem of *Elasmobranchii*.

Our phylogenomic analyses also support post-Cretaceous origins for specialized species of living pelagic sharks and rays (135). The origins of filter feeding in living lineages of sharks and rays is of great interest because, along with the far more recently diversified baleen whales, the four major lineages of planktivorous elasmobranchs are the only other large-bodied filter-feeding marine vertebrates. However, a variety of lineages, including Jurassic-Cretaceous pachyormid fishes (136–138), Cretaceous elasmobranchs with possible stem-lamniform affinities (135), the Devonian ‘placoderm’ stem-gnathostome †*Titanichthys* (139), and Cambrian-Ordovician radiodont stem-arthropods (140, 141) have independently exploited the giant planktivore niche over 500 million years of Earth History. Our time-calibrated phylogeny (Figure 2) places the origination of three (manta rays, *Mobulidae*; Whale Shark, *Rhincodon typus*; Basking Shark, *Cetorhinus maximus*) of the four living lineages of giant, pelagic (Figure 3A) planktivorous chondrichthyans after the Cretaceous-Paleogene boundary and following the extinction of giant filter-feeding fishes and ray-like lamniforms (135, 136); this result is consistent with the fossil record (135–137, 142). We confirm that least two of these lineages, the *Mobula* rays and the Whale Shark, evolved among ancestrally benthopelagic, rather than pelagic, clades, based on ancestral state reconstructions on the time-calibrated phylogeny (Figure 3A). The Cenozoic age of living planktivore sharks and rays inferred using our UCE dataset are congruent with the hypothesis that plankton turnover and extinction at the Cretaceous-Paleogene boundary may have contributed to the faunal turnover of giant filter-feeding vertebrates (135, 136, 143). Although genomic data for Megamouth Shark *Megachasma pelagios* is unavailable, our Bayesian phylogeny of elasmobranchs based on *rag1* sequences, which places this taxon sister to sand sharks (*Odontaspis ferox*), supports the inference (144, 145) that this species independently acquired filter feeding.

**Figure 3.**
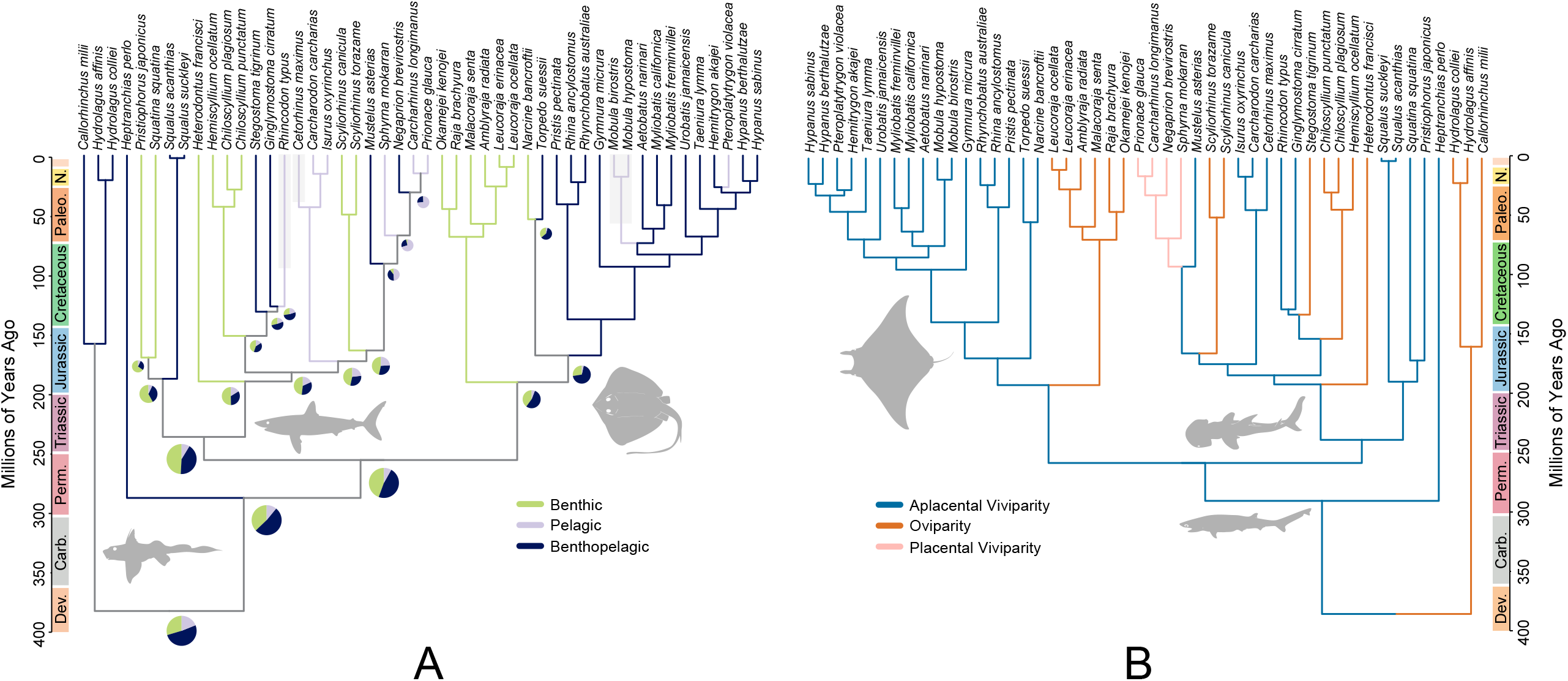
Evolution of chondrichthyan ecology and life history. Phylogenetic ancestral state reconstructions of (A) ecological category and (B) parity mode along the time-calibrated phylogeny of *Chondrichthyes* generated using 356 ultraconserved elements. Illustrations are public domain.

### Ecology, life history, and genomic evolution of sharks and rays

*Elasmobranchii* is notable for the apparent convergent evolution of multiple modes of viviparity within the clade (42, 47, 146–149). Our ancestral state reconstructions of parity mode using the phylogeny of *Chondrichthyes* inferred using UCEs (Figure 3B) indicates a higher degree of uncertainty associated with the ancestral parity mode of cartilaginous fishes than inferred in previous studies, which reconstruct oviparity as the ancestral state (42, 150–152). As in previous studies, we find a high degree of uncertainty underlies inferred ancestral parity states in sharks (42, 150), which is likely driven by the presence of both oviparous and viviparous species in several orders (Figure 3B). Indeed, analyses of larger taxonomic samples of chondrichthyans using phylogenies inferred from molecular sequence data have generally found that oviparity precedes viviparity and reversions to oviparity are rare (150–152). Given that multiple lineages of stem-gnathostome ‘placoderms’(153, 154) and some stem-holocephalans (155) were viviparous, our study raises the intriguing possibility of complex parity mode evolution proximal to the common ancestor of crown gnathostomes (Figure 3B). Notably, we also find that all planktivorous lineages ancestrally possessed aplacental viviparity (Figure 3B) (42, 61). Given recent interest in how paedomorphosis (156) and increased maternal investment (42, 151, 156) might be associated with adaptations for filter-feeding and body size increases, respectively, in sharks and rays, our findings are intriguing for supporting the existence of a common developmental mode in these large pelagic filter-feeding lineages.

Our understanding of chondrichthyan genome evolution has been hampered by the massive genomes of elasmobranchs (46) which outside of lungfishes (89, 90) and lissamphibians (157–160) are the largest known vertebrate genomes (43, 47, 48). The evolution of genome size in sharks, skates, rays, and other vertebrates has been related to life history and metabolic evolution (47, 158, 160). Using measurements of published genome sequence lengths (ln-gigabase pairs) and genome weight (47), we reconstructed ancestral genome size across *Chondrichthyes* to infer the timescale of genomic expansion in this clade. Our reconstruction of genome size evolution infers a single increase in genome size at the common ancestor of *Elasmobranchii* (Figure 4A) and implies high subclade disparity in genome size characterized early crown chondrichthyan evolution (Figure 4B). Observed disparity in genome size from the start of the Mesozoic to the present is not outside the expectations of Brownian motion (Figure 4B). A recent study that examined shark genome size evolution (as measured in picograms) found evidence for an initial increase in genome size at the common ancestor of sharks (monophyletic in their reference tree) followed by substantial, parallel expansions in *Heterodontus* and all squalomorphs except *Hexanchiformes* [figure 1 in (47)]. In contrast, ancestral state reconstructions suggest that the comparatively small genome sizes of hexanchiform sharks reflect their early divergence from other sharks, rays, and skates, rather than being a secondary reduction (Figure 4A; Figure S9).

**Figure 4.**
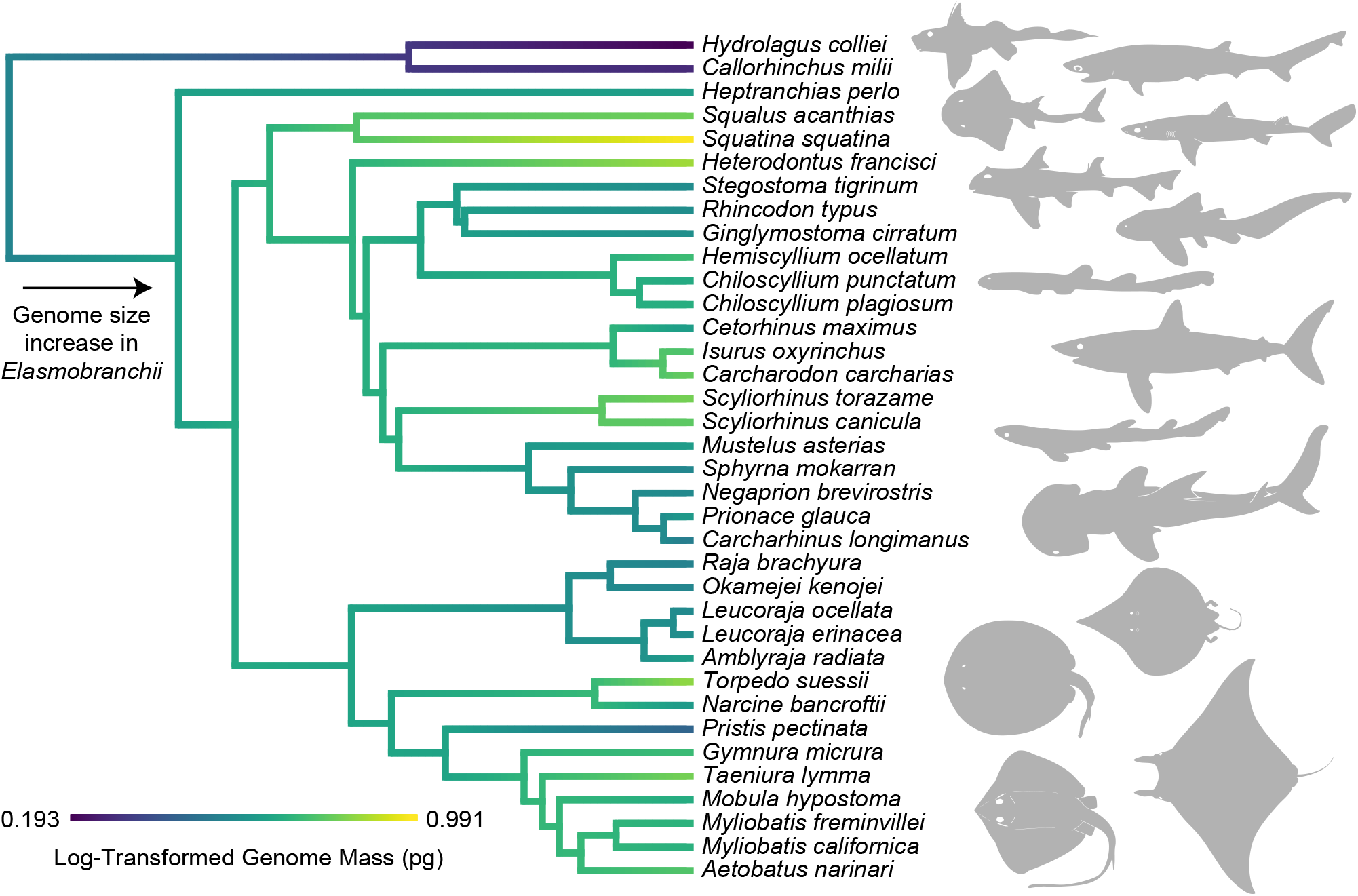
Evolution of chondrichthyan genome size. Traitgram of natural-log-transformed genome weight in picograms made along the time-calibrated phylogeny of *Chondrichthyes* generated using 356 ultraconserved elements. Illustrations are public domain.

## Conclusions

Using newly-assembled datasets of UCEs and BUSCOs from 38 chondrichthyan genomes spanning the major shark, ray, and skate clades, we are able to resolve shark relationships with the exception of the placement of *Hexanchiformes*, which is placed alternatively as the sister to all other shark lineages or as the deepest divergence in *Squalomorphii* depending on the dataset used (Figure 1, Figure 2). Although the resoluiton of this lineage is a hard phylogenetic problem, our analyses suggest that analyses of BUSCOs, which are exons, might be affected by compositional biases. Previous studies of avian (66, 71), cichlid (79), and frog (161) phylogenetics have also found that UCEs outperform exons and are more robust to data subsampling strategy. The phylogenomic hypothesis of shark evolution wherein traditional sharks are paraphyletic (Figure 1, Figure 2) suggests that the cranial anatomy and genome sizes of *Hexanchiformes* are reflective of their deeply divergent phylogenetic position rather than secondary reversions following modifications to jaw articulation and genome size expansion at the base of *Elasmobranchii* (47, 86, 87). Our time-calibrated phylogeny supports a Jurassic diversification of most sharks and a Cretaceous-Paleogene diversification of rays, skates, and relatives (Figure 2). We also infer post-Cretaceous origins of living giant planktivore chondrichthyans (Figure 2; Figure 3A), supporting the hypothesis that these taxa occupied that niche following the extinction of Mesozoic filter-feeding teleosts and chondrichthyans (135, 136). By clarifying where phylogenetic uncertainty remains in the relationships of one of the two major divisions of crown-group jawed vertebrates, we illuminate the unexpectedly ancient evolutionary history of some chondrichthyans that are now under threat of extinction due to the current anthropogenic biodiversity crisis (32).

## Methods

### Taxonomy

In this manuscript, we follow the taxonomic conventions of phylogenetic rank-free taxonomy formalized in the PhyloCode (162), which has recently been applied to both ray-finned fishes (*Actinopterygii*)(163–165) and chimaeras and ratfishes (*Holocephali*)(166). In practice, this means following emerging conventions for italicizing clade names (167), using the prefix pan-to refer to total clades and stem lineages of crown clades, and finally removing redundant clade names. In this manuscript, for example, we do not use the traditional crown ‘orders’ for Bramble Sharks (*Echinorhiniformes*) and Angel Sharks (*Squatiniformes*), since each of these clades is represented by one living genus each (*Echinorhinus* and *Squatina*). Thus, *Echinorhiniformes* is redundant with *Echinorhinus*, and *Squatiniformes* with *Squatina* according to the PhyloCode.

### UCE Sequence Dataset Assembly

In order to leverage genome-scale data to analyze chondrichthyan relationships, we used phyluce 1.7.3 (168) to extract ultraconserved elements from 48 published chondrichthyan genomes and six outgroups, align them, and produce 75% and 90% complete matrices to assess the effect of data completeness on inference of shark interrelationships. We used the 5k vertebrate ultraconserved element probe set (59). Following UCE extraction and alignment creation, we used CIAlign (169) to visualize and prune chimaeric UCE sequence alignments. Our final set consisted of 356 ultraconserved element sequences for 48 chondrichthyans and six osteichthyan outgroups: *Homo sapiens, Latimeria chalumnae, Polypterus senegalus, Scleropages formosus, Amia ocellicauda* (labeled *Amia calva* on Genbank; S. David pers. comm.), and *Lepisosteus oculatus*. We conducted all UCE dataset assembly steps on the Yale high-performance computing cluster McCleary.

### BUSCO Sequence Dataset Assembly

In order to assess how different genomic datasets might affect inference of chondrichthyan relationships (170), we used Augustus in BUSCO 5.8.3 (171, 172) to locate and extract 954 genes conserved across metazoans using the metazoa_odb10 dataset, following a previous study (49). Several genomes were not of sufficient quality for BUSCO genes to be found, so only a total of 45 genomes were used. We collected and aligned the output nucleotide sequences using MAFFT (173) and then used commands in phyluce-1.7.3 and CIAlign to reduce the alignment to 75% and 90% complete matrices and filter chimaeric alignments. Our final 75% complete set consisted of 840 BUSCO sequences for 39 chondrichthyans and 6 osteichthyan outgroups, and our 90% complete set consisted of 483 BUSCO sequences for the same species. We conducted all BUSCO dataset assembly steps on the Yale high-performance computing cluster McCleary.

### Legacy Nuclear Gene Dataset Assembly and Analysis

To exploit the availability of legacy nuclear gene sequence data for a larger sample of chondrichthyans, we aligned all sequences of recombination activating gene 1 (*rag1*) available for 161 chondrichthyans on the NCBI repository GenBank, then ran an uncalibrated Bayesian phylogenetic analysis using MrBayes 3.2.7 (174). For this analysis, we partitioned the sequences by codon position, used an HKY+I+G model, and ran four chains over 10 million generations sampling every thousand generations. Finally, we checked for convergence of the posteriors using Tracer 1.7.1 (175) and summarized the posterior tree sets in a single maximum clade credibility tree.

### Maximum Likelihood Phylogenetic Analysis of Genomic Data

We conducted maximum likelihood phylogenetic analyses in IQ-TREE2 (176, 177) on the concatenated 75% and 90% UCE and BUSCO sequence datasets. Concatenated datasets were alternately treated as a single partition or multiple partitions; in the latter case, we used PartitionFinder 2 (178) to select partitioning schemes. Gene trees were inferred from each UCE locus using IQ-TREE2, models of nucleotide evolution were chosen using ModelFinder as implemented in IQ-TREE2 (179). Node support was assessed using 1000 ultrafast bootstrap replicates. We used the individual gene trees to generate a species tree using the coalescent model implemented in ASTRAL-III (180). Next, we explored site-wide and gene-wide nodal support for the inferred phylogenies using gene and site concordance factors, which measure the number of decisive gene trees and sites that support a given topology in an input tree (here, the single partition concatenated phylogeny) (181). Using custom scripts in ggplot2 (https://www.robertlanfear.com/blog/files/concordance_factors.html), we plotted concordance factors against one another and branch lengths in the maximum likelihood phylogeny. Next, we used commands in phyluce-1.7.3 to collect information on nucleotide base content and parsimony informative site counts for both the BUSCO and UCE datasets at varying levels of completeness. All phylogenetic analyses were conducted on the Yale high-performance computing cluster McCleary.

### Bayesian Time-Calibration

We inferred a time-calibrated phylogeny of *Chondrichthyes* using a Bayesian node-dating approach implemented in BEAST 2.6.6 (182, 183) for three sets of 50 UCEs (owing to their higher taxon sampling) from the 90% complete matrix. We applied a stringent set of criteria to choose a set of 14 fossil calibrations (53, 105) to calibrate the phylogeny, as the fossil record of chondrichthyans includes numerous occurrences based on fragmentary or isolated fossils that may simply represent species that converged, for example, on the dental anatomy of living species. As such, we searched for holomorphic fossils that are universally agreed to represent members of the relevant crown clades and have been placed in phylogenetic analyses of morphological characters (Supplementary Text). The final fossil calibration dataset included 12 species based on holomorphic specimens and two tooth taxa commonly used in time calibration of chondrichthyan phylogeny (63, 105).

For all three UCE sets, we constructed input xml files in BEAUTi 2.6.7 (182, 183). We used a general-time-reversable model of molecular sequence evolution with a gamma site model, a relaxed log-normal clock model, and the fossilized birth-death branching model as implemented in BEAST2 (184). We set the origin prior to 443.8 Ma, the Ordovician-Silurian boundary, as crown gnathostomes are known up to, but not before, the start of the Silurian (31, 31, 185). The bounds on the origin priors were 439.0 Ma, the age of the oldest anatomically well-characterized crown-group gnathostome (185), and 509.0, the minimum age of the stem-vertebrate-bearing (186) Burgess-Shale. We set the diversification rate to 0.018, which is the mean diversification rate estimate according to the formula of (187) given 1489 living chondrichthyan species (188) and an origin of 443.8 Ma, with bounds of 0.00 and 1.00. Next, we set rho, the proportion of living species sampled, to 0.032 (48/1489). We constrained the topology to the ASTRAL-III phylogeny generated from analysis of the 90% complete UCE matrix, ran three independent BEAST2 chains over 600 million generations with a 50 million generation pre-burnin on the Yale high performance computing cluster McCleary, and checked for convergence of the posteriors and effective sample size values over 200 in Tracer v. 1.7.1 (175). After combining the 9 tree sets in LogCombiner 2.6.7 (182) with 75% burnin sampling every 5000 generations, we annotated the ASTRAL-III target tree with median node height values in TreeAnnotator v. 2.6.6 (182).

### Ancestral State Reconstructions and Disparity Through Time

We compiled data on shark habitat preference (pelagic/benthopelagic/benthic)(61) and parity mode from Fishbase and previous studies (42, 150) and conducted ancestral state reconstructions on the Bayesian time-calibrated phylogeny using stochastic mapping over 1000 simulations using the R package phytools (189, 190). For our analyses of genome sizes, we collected data on genome sequence length in gigabase pairs and chromosome number from Genbank using given GenBank lengths and data on genome size in picograms from a previous study (47) and the Animal Genome Size Database (https://www.genomesize.com/; chromosome numbers were also collected from this database if needed). If other species in a genus included in our phylogeny were represented in the Animal Genome Size Database, we took the mean genome size value for these species as a proxy for genome size of the species included in our tree. In one case, we reduced our sample of *Mobula* to a single tip to institute this approach. Because no data is available on total genome size in picograms for *Heptrachias perlo*, we took the average of genome sizes reported for members of *Hexanchiformes* and inputted the value for that taxon, which represents this whole clade in our tree. Next, we used scripts in the R packages phytools and geiger (191) to fit four models of evolution (Brownian Motion, Ornstein-Uhlenbeck, White Noise, and Early Burst) on the log-transformed genome sequence length and size data, produce traitgrams scaled by these measurements, and produce disparity-through-time plots of genome length and size over time.

## Acknowledgements

We thank members of the Near Lab for discussions related to this manuscript. We also thank the editor and reviewers for their helpful comments.

