## Supplementary material for "Phylogenomics and the origins of sharks": Table 1

Table 1. Counts of parsimony informative sites for each analysis.

|  | Analysis | Mean | Min | Max | Total |
| --- | --- | --- | --- | --- | --- |
|  | UCEs, 75% | 362.44 | 59 | 596 | 126490 |
|  | UCEs, 90% | 370.61 | 59 | 546 | 75976 |
|  | BUSCOs, 75% | 652.77 | 85 | 4548 | 548324 |
|  | BUSCOs, 90% | 656.3 | 124 | 4548 | 316991 |
