## Supplemental Text and Tables for "Phylogenomics and the origins of sharks"

**Supplementary Text: Phylogenomics and the origins of sharks**

Chase D. Brownstein,[1*] and Thomas J. Near[1,2]
[1]Department of Ecology and Evolutionary Biology, Yale University, New Haven CT, USA
[2]Yale Peabody Museum, New Haven CT, USA

**This file includes:**
Fossil Calibration Justification List

**Fossil Calibration Justification List.**
For the purposes of this study, we follow best practices of fossil calibration justification outlined in Parham et al. (*1*). Below, we provide justifications for the position of fossils based on comparative anatomy and phylogenetically optimized morphological character states. Chondrichthyan workers often rely on the excellent shark tooth record to understand the timing of origination of families and genera (*2–5*). We have attempted to use as many holomorphic fossils as possible to calibrate our phylogeny simply because holomorphic fossils preserve more phylogenetically informative character data, and nearly all of these calibrations are equal in age to the earliest unambiguous tooth records of clades. However, when we choose to use a holomorphic fossil that postdates the age of a tooth that is identifiable to the same taxon as a younger holomorphic fossil, we explicitly note so. A total of four of our 16 fossil calibrations are based on teeth; all of the tooth taxa used have previously been vetted in comprehensive node calibration justifications (*3*, *6*)

†*Guiyu oneiros*
**Justification of Placement:** †*Guiyu oneiros* calibrates the MRCA of crown *Osteichthyes* (specifiers: *Homo sapiens*, *Amia calva*) in our node-dating analysis. The placement of †*Guiyu oneiros* within crown *Osteichthyes* is supported by multiple parsimony and Bayesian phylogenetic analyses of morphological characters, including the 153 character, 23 taxon matrix of Zhu et al. (*7*), the 236 character, 78 taxon matrix of Giles et al. (*8*), the 182 character, 49 taxon matrix of Giles et al. (*9*), the 269 character, 96 taxon matrix of Lu et al. (*10*), the 335 character, 103 taxon matrix of Qiao et al. (*11*), the 336 character, 104 taxon matrix of Choo et al. (*12*), the 347 character, 104 taxon matrix of Cui et al. (*13*), and the 694 character, 159 taxon matrix of Cui et al. (*14*). Notably, Lu et al. (*15*), King et al. (*16*), and Brazeau et al. (*17*) placed †*G. oneiros* within a clade of Devonian osteichthyans that is the sister lineage to crown *Osteichthyes*. However, analyses of an expanded version of the King et al. (*16*), and Brazeau et al. (*17*) matrices either place this clade in a polytomy with crown osteichthyans or on the sarcopterygian stem under a Bayesian tip-dating approach (*18*). For the purposes of this justification, we rely on the phylogeny presented in supplementary figure 6 in Cui et al. (*14*) Because no character state optimizations were provided in Cui et al. (*14*), we reran the matrix presented in the paper using TNT v 1.5 (*19*) to optimize apomorphies; we ran an initial Wagner search over 1000 replicates and default parameters for ratchet, tfuse, drift, and sectorial search followed by traditional bisection reconnection branch swapping over 100,000 trees. We summarized the topologies in a strict consensus and then recorded apomorphies of two nodes: the clade containing †*G. oneiros* and crown *Osteichthyes* and †*Guiyu oneiros* and Pan-*Sarcopterygii*. Phylogenetically optimized apomorphies uniting †*G. oneiros* with crown *Osteichthyes* are: presence of differentiated lepidotrichia (28:1 in Cui et al. (*14*)), posterior end of supraorbital canal contained in postparietal (178:0 in Cui et al. (*14*)), and posterior expansion of the maxilla (270:0 in Cui et al. (*14*)). Phylogenetically optimized apomorphies uniting †*G. oneiros* with Pan-*Sarcopterygii* are: dermal cranial joint at the level of sphenoid-otic junction (111:1 in Cui et al. (*14*)), presence of the supraorbital

(125:1 in Cui et al. (*14*)), presence of an endoskeletal intracranial joint (371:1 in Cui et al. (*14*)), internasal vacuities shallow, paired pits with strong midline ridge (390:1 in Cui et al. (*14*)), and presence of an unconstricted cranial notochord (411:1 in Cui et al. (*14*)).
**Stratigraphic Horizon**: Qujing, Kuanti Formation, Yunnan Province, China; late Ludfordian Stage of the Silurian (*20*). The Ludfordian Stage ranges between 425.6 and 423.0 Ma (*21*).
**Equivalent Fossil Calibrations:** None. Slightly younger calibrations of the same node include the 'psarolepids' †*Achoania jarvikii* (*22*) and †*Psarolepis romeri* (*23*), as well as the stem actinopterygian †*Meemania eos* (*10*). However, these taxa are younger than the age of the oldest stem-lungfishes (see below)(*24*) and are therefore redundant for fossil calibration.
**Fossil tip age**: 423.0 Ma. The node calibration was created such that 97.5% of the probability distribution fell before 423.0 Ma.

†*Youngolepis praecursor*
**Justification of Placement:** †*Youngolepis praecursor* calibrates the MRCA of *Sarcopterygii* (specifiers: *Homo sapiens*, *Latimeria chalumnae*) in our node dating analyses. The placement of †*Y. praecursor* in *Sarcopterygii* as a member of Pan-*Dipnoi* (lungfishes) is supported by parsimony and Bayesian phylogenetic analyses of morphological characters, including the 335 character, 103 taxon matrix of Qiao et al. (*11*), the 242 character, 37 taxon matrix of Lu et al. (*25*), the 70 character, 25 taxon matrix of Friedman (*26*), and the 277 character, 88 taxon matrix of Cui et al. (*24*). For the purposes of this justification, we rely on the phylogenies presented in supplementary figure 2 in Cui et al.(*24*) Because no character state optimizations were provided by Cui et al. (*24*) for the placement of †*Y. praecursor* sister to all other members of Pan-*Dipnoi*, we reran the matrix presented in the paper using TNT v 1.5 (*19*) to optimize apomorphies; we ran an initial Wagner search over 1000 replicates and default parameters for ratchet, tfuse, drift, and sectorial search followed by traditional bisection reconnection branch swapping over 100,000 trees. We summarized the topologies in a strict consensus and then recorded apomorphies of †*Y. praecursor*+Pan-*Dipnoi.* Phylogenetically optimized apomorphies uniting †*Y. praecursor* with Pan-*Dipnoi* are: closed pineal opening (1:1 in Cui et al. (*24*)), quadratojugal as large as jugal (34:0 in Cui et al. (*24*)), adductor fossa more than one third the length of the jaw (133:0 in Cui et al. (*24*)), addition of large dentine elements at regular intervals to lateral margin of pterygoid and/or prearticular (166:0 in Cui et al. (*24*)), dentine sheet on resorption areas within tooth plate on palate (174:1 in Cui et al. (*24*)), marginal tooth ridge continuous (185:1 in Cui et al. (*24*)), intracranial joint present as a ventral cranial fissure (190:1 in Cui et al. (*24*)), and a triangular hyomandibular (271:1 in Cui et al. (*24*)).
**Stratigraphic Horizon**: Yunnan, China; Xitun Formation, earliest Lochkovian Stage of the Devonian (*24*, *27*). We followed Cui et al. (*24*) as treating its age as 418.0 Ma.
**Equivalent Fossil Calibrations:** †*Diabolepis speratus* is known from the same formation and is unambiguously placed as a stem-dipnoan crownward of †*Youngolepis praecursor* in the phylogeny of Cui et al. (*24*), as well as a pan-dipnoan in other phylogenetic analyses (*11*, *17*, *28–30*).
**Fossil tip age**: 418.0 Ma. The node calibration was created such that 97.5% of the probability distribution fell before 418.0 Ma.

†*Watsonulus eugnathoides*
**Justification of Placement:** †*Watsonulus eugnathoides* calibrates the MRCA of *Holostei* (specifiers: *Amia calva*, *Lepisosteus oculatus*) in our node-dating analyses. The placement of †*W. eugnathoides* in *Holostei* as a member of Pan-*Amia* is supported by Bayesian and parsimony analyses of morphological characters, including the 90 character, 37 taxon matrix of López-Arbarello (*31*), the 96 character, 28 taxon matrix of Xu et al. (*32*), the 265

character, 93 taxon matrix of Giles et al. (*33*), the 339 character, 99 taxon matrix of López-Arbarello and Sferco (*34*), the 275 character, 97 taxon dataset of Argyriou et al. (*35*), the 224 character, 60 taxon dataset of Xu (*36*), and the 300 character, 117 taxon matrix of Argyriou et al. (*37*). For the purposes of this justification, we rely on the phylogeny presented in figure 11 of Argyriou et al. (*37*). Phylogenetically optimized apomorphies uniting †*W. eugnathoides* with Pan-*Amia* are: posterior maxillary notch present (75:1 in Argyriou et al. (*37*)), single supramaxilla (76:1 in Argyriou et al. (*37*)), accessory hyoid element involved in jaw joint (97:1 in Argyriou et al. (*37*)), foramen for abducens nerve (VI) level with optic foramen (II) (135:1 in Argyriou et al. (*37*)), anterior ossification of ceratohyal unconstructed (220:0), and jaw articulation posterior to orbit (280:0 in Argyriou et al. (*37*)).
**Stratigraphic Horizon**: Ambilombe, Madagascar; Sakamena Formation, Early Triassic, Induan (*38*). We use an age of 251.2 Ma, which is the base of the Induan (*21*), following previous studies (*39*).
**Equivalent Fossil Calibrations:** None.
**Fossil tip age**: 251.2 Ma. The node calibration was created such that 97.5% of the probability distribution fell before 251.2 Ma.

†*Phoebodus saidselachus*
**Justification of Placement:** †*Phoebodus saidselachus* calibrates the most recent common ancestor (MRCA) of crown *Chondrichthyes* (specifiers: *Callorhinchus milii*, *Carcharodon carcharias*) in our node-dating analysis. The placement of †*P. saidselachus* within crown *Chondrichthyes* on the stem of *Elasmobranchii* (sharks, skates and rays) is supported by parsimony and Bayesian phylogenetic analyses of morphological characters, including the 221 character, 60 taxon matrix of Frey et al. (*40*), the 230 character, 64 taxon matrix of Frey et al. (*41*, *42*), and the 236 character, 36 taxon matrix of Brownstein et al. (*43*). All of these matrices are modified from the matrix of Coates et al. (*44*). For the purposes of this justification, we rely on the phylogeny presented in figure 4 in Frey et al. (*40*) as the reference tree. Phylogenetically optimized apomorphies uniting †*P. saidselachus* with Pan-*Elasmobranchii* are: smooth ceratohyal has a posterior, lateral fossa (character 51:1 in Frey et al. (*40*)), dorsal otic ridge forms a crest posteriorly (156:1 in Frey et al. (*40*)), slot-shaped endolymphatic fossa divides dorsal otic ridge along midline (158:1 in Frey et al. (*40*)), dorsal portion of occipital arch wedged between otic capsules (171:1 in Frey et al. (*40*)), posterior/pelvic-level dorsal fin possesses calcified base plate (200:1 in Frey et al. (*40*)).
**Stratigraphic Horizon**: Madene El Mrakib and Aguelmous, Ibâouane Formation (Lahfira Member, Thylacocephalan Layer)(*41*), southern Maïder region, eastern Anti-Atlas, Morocco; Middle Famennian Stage of the Devonian (*40*), 372.2 to 359.3-358.9 Ma (*21*, *45*).
**Equivalent Fossil Calibrations:** †*Ferromirum oukherbouchi* and †*Maghriboselache mohamezanei* are chondrichthyans known from whole-body fossils from the same stratigraphic horizon that are unambiguously placed as stem-holocephalans in crown *Chondrichthyes* in parsimony and Bayesian phylogenetic analyses of morphological characters (*41–43*).
**Fossil tip age**: 358.9 Ma. The node calibration was created such that 97.5% of the probability distribution fell before 358.9 Ma.

†*Ischyodus quenstadti*
**Placement**: †*Ischyodus quenstadti* calibrates the most recent common ancestor of *Holocephali* (specifiers: *Callorhinchus milii, Chimaera colliei*) in our node-dating analysis. The placement of †*Ischyodus quenstadti* as a member of Pan-*Callorhinchus* is supported by parsimony and Bayesian phylogenetic analyses of morphological characters, including the 236 character, 36 taxon matrix of Brownstein et al. (*43*). For the purposes of this justification,

we rely on the phylogeny presented in figure 1 of Brownstein et al. (*43*) as the reference tree. Because no character state optimizations were provided in Brownstein et al. (*43*), we reran the matrix presented in the paper using TNT v 1.5 (*19*) to optimize apomorphies; we ran an initial Wagner search over 1000 replicates and default parameters for ratchet, tfuse, drift, and sectorial search followed by traditional bisection reconnection branch swapping over 100,000 trees. We summarized the topologies in a strict consensus and then recorded apomorphies of *Holocephali* and Pan-*Callorhinchus*. Phylogenetically optimized apomorphies uniting †*I. quenstadti* with *Holocephali* are: hypermineralised regions on all toothplates (tritors) present on all toothplates (81:1 in Brownstein et al. (*43*)). Phylogenetically optimized apomorphies uniting †*I. quenstadti* with Pan-*Callorhinchus* are: tritoral dentine not organized into rod-like structures (82:0 in Brownstein et al. (*43*)), toothplates with prominent descending lamina (83:1in Brownstein et al. (*43*)), rostral cartilage supports plough-shaped snout (108:1 in Brownstein et al. (*43*)), anal fin present (221:1 in Brownstein et al. (*43*)), caudal fin heterocercal in shape (225:0 in Brownstein et al. (*43*)).
**Stratigraphy**: Various localities, Solnhofen Plattenkalk, Altmühltal Formation, Solnhofen, Germany; Tithonian (*46*), 149.2 to 145.0 Ma (*21*).
**Equivalent Fossil Calibrations:** †*Elasmodectes* sp. from the Solnhofen Plattenkalk, as well as isolated egg cases potentially referable to crown holocephalans from the same unit, are equivalent fossil calibrations (*43*, *46*).
**Fossil tip age**: 145.0 Ma. The node calibration was created such that 97.5% of the probability distribution fell before 145.0 Ma.

†*Heterodontus zitteli*
**Placement**: †*Heterodontus zitteli* calibrates the most recent common ancestor of *Galeomorphii* (specifiers: *Heterodontus francisci*, *Carcharodon carcharias*) in our node-dating analysis. The placement of †*H. zitteli* in *Galeomorphii* as a member of Pan-*Heterodontus* is supported by parsimony phylogenetic analysis of morphological characters, including the 113 character, 10 taxon matrix of Slater et al. (*47*). For the purposes of this justification, we rely on the phylogeny presented in figure 6 of Slater et al. (*47*) as the reference tree. Phylogenetically optimized apomorphies uniting of †*H. zitteli* with Pan-*Heterodontus* are: low number of tooth families, absence of labial tooth crown ornamentation on anterior teeth, anal fin that is more than its own length in distance to the caudal fin, pectoral girdle positioned at the eighth vertebrae (*47*).
**Stratigraphy:** Various localities, holotype is from Eichstaett Plattenkalk, Germany; Kimmeridgean-Tithonian, 149.2 Ma (*46*, *47*).
**Equivalent Fossil Calibrations:** Various; other Pan-*Heterodontus* of equivalent age include †*Paracestracion* spp. (*47*).
**Fossil tip age**: 149.2 Ma. The node calibration was created such that 97.5% of the probability distribution fell before 149.2 Ma.

†*Pseudorhina* sp.
**Placement**: †*Pseudorhina* sp. calibrates the most recent common ancestor of *Squatina squatina* and *Pristiophorus japonicus* in our node-dating analysis. The placement of †*P.* spp. as a member of Pan-*Squatina* is supported by parsimony and maximum likelihood analyses of morphological characters, including the 224 character, 52 taxon matrix of Jambura et al. (*48*) and the 221 character, 91 taxon matrix of Vullo et al. (*49*). For the purposes of this justification, we rely on the phylogeny presented in figure 9 of Jambura et al. (*48*). Phylogenetically optimized apomorphies uniting †*Pseudorhina* with Pan-*Squatina* are: nasal capsules anteriorly expanded (15:2 in Jambura et al. (*48*)), posterior margin of puboischiadic bar posteriorly directed (119:1 in Jambura et al. (*48*)), tearing dentition (183:1 in Jambura et

al. (*48*)), arched mesial and distal tooth crowns present (201:1 in Jambura et al. (*48*)), lower teeth apron narrow, high extended below half the height of the root, thick, conical, and uniform overhanging the basal face of the root (222:4 in Jambura et al. (*48*)).
**Stratigraphy:** Various localities. The oldest holomorphic fossil of †*Pseudorhina acanthoderma* is from the late Kimmeridgian (*Hybonoticeras beckeri* Zone, *Lithacoceras ulmense* Subzone) of Nusplingen, Germany (*5*). Teeth unambiguously belonging to †*Pseudorhina* sp. have been recovered from units as old as the upper Oxfordian, 156.2 Ma (*5*).
**Equivalent Fossil Calibrations:** †*Protospinax* spp. from the Early to Late Jurassic of Europe is a potentially equivalent fossil calibration (*48*); however, owing to disagreement about the placement of this taxon among sharks (*48*, *50*), we use the unambigious (*5*, *46*, *48*, *48*, *51*) Pan-*Squatina* †*Pseudorhina acanthoderma* instead.
**Fossil tip age**: 156.2 Ma. The node calibration was created such that 97.5% of the probability distribution fell before 156.2 Ma.

†*Cantioscyllium alhaulfi*
**Placement**: †*Cantioscyllium alhaulfi* calibrates the most recent common ancestor of *Ginglymostoma cirratum* and *Rhincodon typus* in our node-dating analysis. The placement of †*C. alhaulfi* as a member of Pan-*Ginglymostomatidae* is supported by characteristics of the dentition (*52*). Previous studies have also employed this taxon as a fossil calibration (*3*, *6*).
**Stratigraphy:** Limestones at Alcaine, Teruel Spain; Barremian, 121.4 Ma (*53*).
**Equivalent Fossil Calibrations:** None.
**Fossil tip age**: 121.4 Ma. The node calibration was created such that 97.5% of the probability distribution fell before 121.4 Ma.

†*Palaeoscyllium tenuidens*
**Placement**: †*Palaeoscyllium tenuidens* calibrates the most recent common ancestor of *Carcharhiniformes* (specifiers: *Scyliorhinus canicula, Prionace glauca*) in our node-dating analyses. The placement of †*Palaeoscyllium* spp. as a member of Pan-*Scyliorhinidae* is supported by characteristics of the dentition, including the presence of delicate, high, and strongly ornamented central and lateral cusps (*54*). Previous studies have also employed this taxon as a fossil calibration (*3*, *6*).
**Stratigraphy:** Various localities, England; Bathonian, 165.3 Ma minimum age (*6*).
**Equivalent Fossil Calibrations:** †*Eypea leesi*, another tooth taxon from the Bathonian of England, is an equivalent fossil calibration (*55*).
**Fossil tip age**: 165.3 Ma. The node calibration was created such that 97.5% of the probability distribution fell before 165.3 Ma.

†*Abdounia africana*
**Placement**: *†Abdounia africana* calibrates the most recent common ancestor of *Sphyrna mokarran* and *Prionace glauca* in our node-dating analysis. The placement of *†A. africana* as a member of Pan-*Carcharhinidae* is supported by characteristics of the dentition (*3*, *56*). Previous studies have also employed this taxon as a fossil calibration (*3*, *6*).
**Stratigraphy:** Various localities, Paleogene; 61.6 Ma (*3*).
**Equivalent Fossil Calibrations:** None.
**Fossil tip age**: 61.6 Ma. The node calibration was created such that 97.5% of the probability distribution fell before 61.6 Ma.

†*Keasius taylori*

**Placement**: †*Keasius taylori* calibrates the most recent common ancestor of *Carcharodon carcharias* and *Cetorhinus maximus* in our node-dating analysis. The placement of †*K. taylori* as a member of Pan-*Cetorhinus* based on characteristic dental and gill raker morphology of this clade (reviewed in detail in (*57*)).
**Stratigraphy:** Various localities, middle member of the Keasey Formation, Columbia County, Oregon; Duchesnean North American land-mammal age, Priabonian Stage, Eocene Epoch, Paleogene, 40.4-39.5 Ma (*57*).
**Equivalent Fossil Calibrations:** None.
**Fossil tip age:** 39.5 Ma. The node calibration was created such that 97.5% of the probability distribution fell before 39.5 Ma.

†"*Dasyatis*" *speetonensis*
**Placement**: †"*Dasyatis*" *speetonensis* calibrates the most recent common ancestor of *Mobula birostris* and *Pristis pectinata* in our node-dating analysis. The placement of †"*Dasyatis*" *speetonensis* as a member of Pan-*Myliobatiformes* is supported by characteristics of its dentition (*58*). This calibration has been used in previous studies (*59*).
**Stratigraphy:** Speeton Clay Formation, northeast England; Hauterivian Stage of the Early Cretaceous (*58*), 132.6-125.77 Ma (*21*).
**Equivalent Fossil Calibrations:** None.
**Fossil tip age:** 125.77 Ma. The node calibration was created such that 97.5% of the probability distribution fell before 125.77 Ma.

†*Titanonarke molini*
**Placement**: †*Titanonarke molini* calibrates the most recent common ancestor of *Narcine bancroftii* and *Torpedo suessii* in our node-dating analyses. The placement of †*T. molini* as a member of *Narcinidae* is supported by parsimony analyses of morphological characters, including the 72 character, 16 taxon matrix of Marramà et al. (*60*). For the purposes of this justification, we rely on the phylogeny presented in figure 18 of Marramà et al. (*60*). Phylogenetically optimized apomorphies uniting †*Titanonarke* with *Torpediniformes* are: electric organs present (1:1 in Marramà et al. (*60*)), dermal denticles and thorns absent (2:1 in Marramà et al. (*60*)), iliac process straight (32:1 in Marramà et al. (*60*)), prepelvic process long (33:1 in Marramà et al. (*60*)), suprascapular antimere fused with visible suture (35:1 in Marramà et al. (*60*)), and basibranchial copular rounded with small caudal point (53:1 in Marramà et al. (*60*)). Phylogenetically optimized apomorphies uniting †*Titanonarke* with *Narcinoidea* are: combined length of small labial cartilages are less than length of Meckel's cartilage (21:1 in Marramà et al. (*60*)), palatoquadrate labiolingually and tapers towards symphysis (23:1 in Marramà et al. (*60*)), lateral stay in posterior third of the synarcual (62:2 in Marramà et al. (*60*)), lateral stay anterior margin describing obtuse angle to axis (64:2 in Marramà et al. (*60*)), and tooth cusp length less than half length of cutting edge (68:1 in Marramà et al. (*60*)). Phylogenetically optimized apomorphies uniting †*Titanonarke* with *Narcinidae* are: rostral fontanelle present (44:1 in Marramà et al. (*60*)), 10 or fewer rib pairs (71:1 in Marramà et al. (*60*)), and rostral cartilage connected to antorbital cartilage through lateral appendix (72:1 in in Marramà et al. (*60*)).
**Stratigraphy:** Monte Bolca, near Verona, Italy: early-late Ypresian, Eocene, Paleogene 48.96 to 48.5 Ma (*63*).
**Equivalent Fossil Calibrations:** †*Eotorpedo* spp., based on isolated teeth from the late Paleocene, is a potential equivalent fossil calibration (*60*).
**Fossil tip age:** 48.5 Ma. The node calibration was created such that 97.5% of the probability distribution fell before 48.5 Ma.

†*Pristis lathami*

**Placement**: †*Pristis lathami* calibrates the most recent common ancestor of *Pristis pectinata* and *Rhina ancylostomus* in our node-dating analysis. The placement of †*P. lathami* as a member of Pan-*Pristis* is supported by the following combination of morphological characters: single pair of elliptical lateral canals on rostrum, rostral teeth firmly attached via alveoli (*64*).

**Stratigraphy:** Various, but we use a body fossil from Lafarge Cement quarry, Harleyville, Dorchester County, South Carolina; Cross Member of the Tupelo Bay Formation, 37 Ma.(*64*)

**Fossil tip age:** 37.0 Ma. The node calibration was created such that 97.5% of the probability distribution fell before 37.0 Ma.

**References.**

21. F. M. Gradstein, J. G. Ogg, M. Schmitz, G. Ogg, *The Geologic Time Scale 2020* (Elsevier Science, Amsterdam, The Netherlands, 2021).

37. T. Argyriou, S. Giles, M. Friedman, A Permian fish reveals widespread distribution of neopterygian-like jaw suspension. *eLife* **11**, e58433.

50. J. Maisey, Phylogenetic relationships of the Late Jurassic shark Protospinax Woodward 1919 (Chondrichthyes: Elasmobranchii). *Mesoz. Fishes Syst. Paleoecol. Verl. Dr Friedrich Pfeil Munich* (1996).

57. B. J. Welton, *A New Archaic Basking Shark (Lamniformes: Cetorhinidae) from the Late Eocene of Western Oregon, USA, and Description of the Dentition, Gill Rakers and Vertebrae of the Recent Basking Shark Cetorhinus Maximus (Gunnerus): Bulletin 58* (New Mexico Museum of Natural History and Science, 2013; https://books.google.com/books?hl=en&lr=&id=5vIoCgAAQBAJ&oi=fnd&pg=PP1&dq=A+NEW+ARCHAIC+BASKING+SHARK+(LAMNIFORMES:+CETORHINIDAE)+FROM+THE+LATE+EOCENE+OF+WESTERN+OREGON,+U.S.A.,+AND+DESCRIPTION+OF+THE+DENTITION,+GILL+RAKERS+AND+VERTEBRAE+OF+THE+RECENT+BASKING+SHARK+CETORHINUS+MAXIMUS+(GUNNERUS)&ots=4q3eITChlR&sig=NWTNaUJE7DwgW5qy-W97tXWq_Rw)vol. 58.

67. Squalean Phylogeny a New Framework of"Squaloid" Sharks and Related Taxa | Semantic Scholar. https://www.semanticscholar.org/paper/Squalean-Phylogeny-a-New-Framework-of%22Squaloid%22-and-Shirai/e4dd3d21d597f4e1696a3070d93455c391bd3afe.

68. Chapter 3 – Higher-Level Elasmobranch Phylogeny, Basal Squaleans, and Paraphyly | Semantic Scholar. https://www.semanticscholar.org/paper/Chapter-3-%E2%80%93-Higher-Level-Elasmobranch-Phylogeny%2C-Carvalho/32cb7cfc3cc063f3e245d19f5f5fc511e6ea2a14.

69. X. Vélez-Zuazo, I. Agnarsson, Shark tales: A molecular species-level phylogeny of sharks (Selachimorpha, Chondrichthyes). *Mol. Phylogenet. Evol.* **58**, 207–217 (2011).

70. J. G. Inoue, M. Miya, K. Lam, B.-H. Tay, J. A. Danks, J. Bell, T. I. Walker, B. Venkatesh, Evolutionary Origin and Phylogeny of the Modern Holocephalans (Chondrichthyes: Chimaeriformes): A Mitogenomic Perspective. *Mol. Biol. Evol.* **27**, 2576–2586 (2010).

71. G. J. P. Naylor, J. N. Caira, K. Jensen, K. A. M. Rosana, N. Straube, and C. Lakner, "Elasmobranch Phylogeny: A Mitochondrial Estimate Based on 595 Species" in *Biology of Sharks and Their Relatives* (CRC Press, ed. 2, 2012).

72. G. J. P. Naylor, J. A. Ryburn, O. Fedrigo, J. A. López, "Phylogenetic Relationships among the Major Lineages of Modern Elasmobranchs" in *Reproductive Biology and Phylogeny of Chondrichthyes* (CRC Press, 2005).

73. C. R. L. Amaral, F. Pereira, D. A. Silva, A. Amorim, E. F. de Carvalho, The mitogenomic phylogeny of the Elasmobranchii (Chondrichthyes). *Mitochondrial DNA Part A* **29**, 867–878 (2018).

**Figure S1.** ASTRAL-III phylogeny generated using the 75% complete UCE matrix. All nodes are supported by coalescent supports of 1.0.

**Figure S2.** IQ-TREE2 single partition phylogeny generated using the concatenated UCEs from the 75% complete matrix. All nodes are supported by ultrafast bootstrap supports of 100 unless indicated.

**Figure S3** IQ-TREE2 multiple partition phylogeny generated using the concatenated UCEs from the 75% complete matrix. All nodes are supported by ultrafast bootstrap supports of 100 unless indicated.

**Figure S4.** ASTRAL-III phylogeny generated using the 90% complete UCE matrix. All nodes are supported by coalescent supports of 1.0.

**Figure S5.** IQ-TREE2 single partition phylogeny generated using the concatenated UCEs from the 90% complete matrix. All nodes are supported by ultrafast bootstrap supports of 100 unless indicated.

**Figure S6.** IQ-TREE2 multiple partition phylogeny generated using the concatenated UCEs from the 90% complete matrix. All nodes are supported by ultrafast bootstrap supports of 100 unless indicated.

**Figure S7.** ASTRAL-III phylogeny generated using the 75% complete BUSCO gene matrix. All nodes are supported by coalescent supports of 1.0.

**Figure S8.** IQ-TREE2 single partition phylogeny generated using the concatenated BUSCO genes from the 75% complete matrix. All nodes are supported by ultrafast bootstrap supports of 100 unless indicated.

**Figure S9.** IQ-TREE2 multiple partition phylogeny generated using the concatenated BUSCO genes from the 75% complete matrix. All nodes are supported by ultrafast bootstrap supports of 100 unless indicated.

**Figure S10.** ASTRAL-III phylogeny generated using the 90% complete BUSCO gene matrix. All nodes are supported by coalescent supports of 1.0.
**Figure S11.** IQ-TREE2 single partition phylogeny generated using the concatenated BUSCO genes from the 90% complete matrix. All nodes are supported by ultrafast bootstrap supports of 100 unless indicated.
**Figure S12.** IQ-TREE2 multiple partition phylogeny generated using the concatenated BUSCO genes from the 90% complete matrix. All nodes are supported by ultrafast bootstrap supports of 100 unless indicated.
**Figure S13.** Relationships between gene and site concordance factors and branch lengths for the 75% and 90% complete BUSCO and UCE matrices.
**Figure S14.** Phylogeny of *Chondrichthyes* inferred using Bayesian analysis of a dataset of available chondrichthyan *rag1* nuclear gene sequences, showing the position of Bramble Shark *Echinorhinus brucus* (one of two species in *Echinorhinus*).
**Figure S15.** Phylogram of log-transformed genome size. Bars are 95% confidence intervals for estimated ancestral genome mass at nodes.

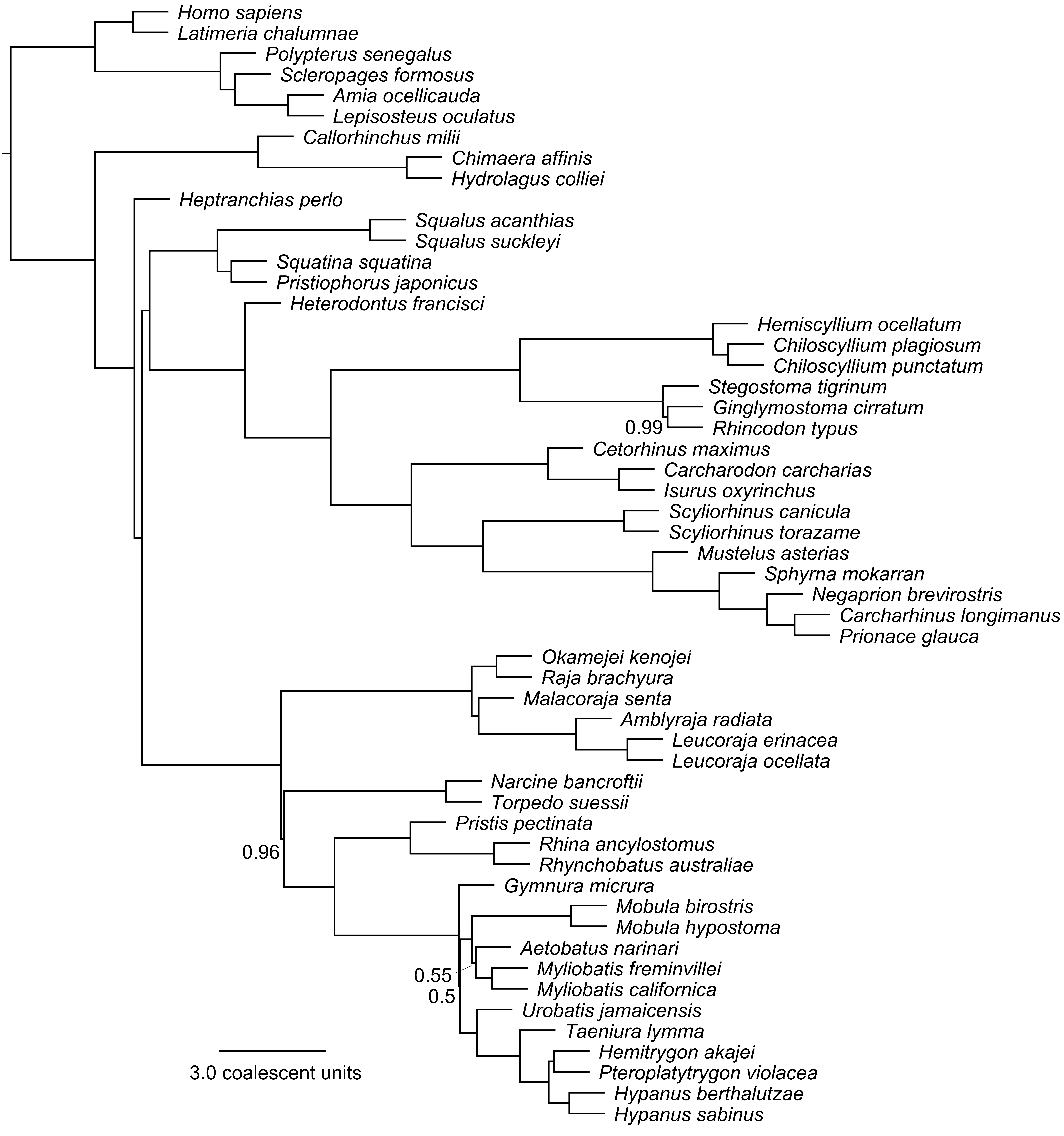
Homo sapiens
Latimeria chalumnae
Polypterus senegalus
Scleropages formosus
Amia ocellicauda
Lepisosteus oculatus
Callorhinchus milii
Chimaera affinis
Hydrolagus colliei
Heptranchias perlo
Squalus acanthias
Squalus suckleyi
Squatina squatina
Pristiophorus japonicus
Heterodontus francisci
Hemiscyllium ocellatum
Chiloscyllium plagiosum
Chiloscyllium punctatum
Stegostoma tigrinum
Ginglymostoma cirratum
Rhincodon typus
0.99
Cetorhinus maximus
Carcharodon carcharias
Isurus oxyrinchus
Scyliorhinus canicula
Scyliorhinus torazame
Mustelus asterias
Sphyrna mokarran
Negaprion brevirostris
Carcharhinus longimanus
Prionace glauca
Okamejei kenojei
Raja brachyura
Malacoraja senta
Amblyraja radiata
Leucoraja erinacea
Leucoraja ocellata
Narcine bancroftii
Torpedo suessii
Pristis pectinata
Rhina ancylostomus
Rhynchobatus australiae
0.96
Gymnura micrura
Mobula birostris
Mobula hypostoma
Aetobatus narinari
Myliobatis freminvillei
Myliobatis californica
0.55
0.5
Urobatis jamaicensis
Taeniura lymma
Hemitrygon akajei
Pteroplatytrygon violacea
Hypanus berthalutzae
Hypanus sabinus
3.0 coalescent units

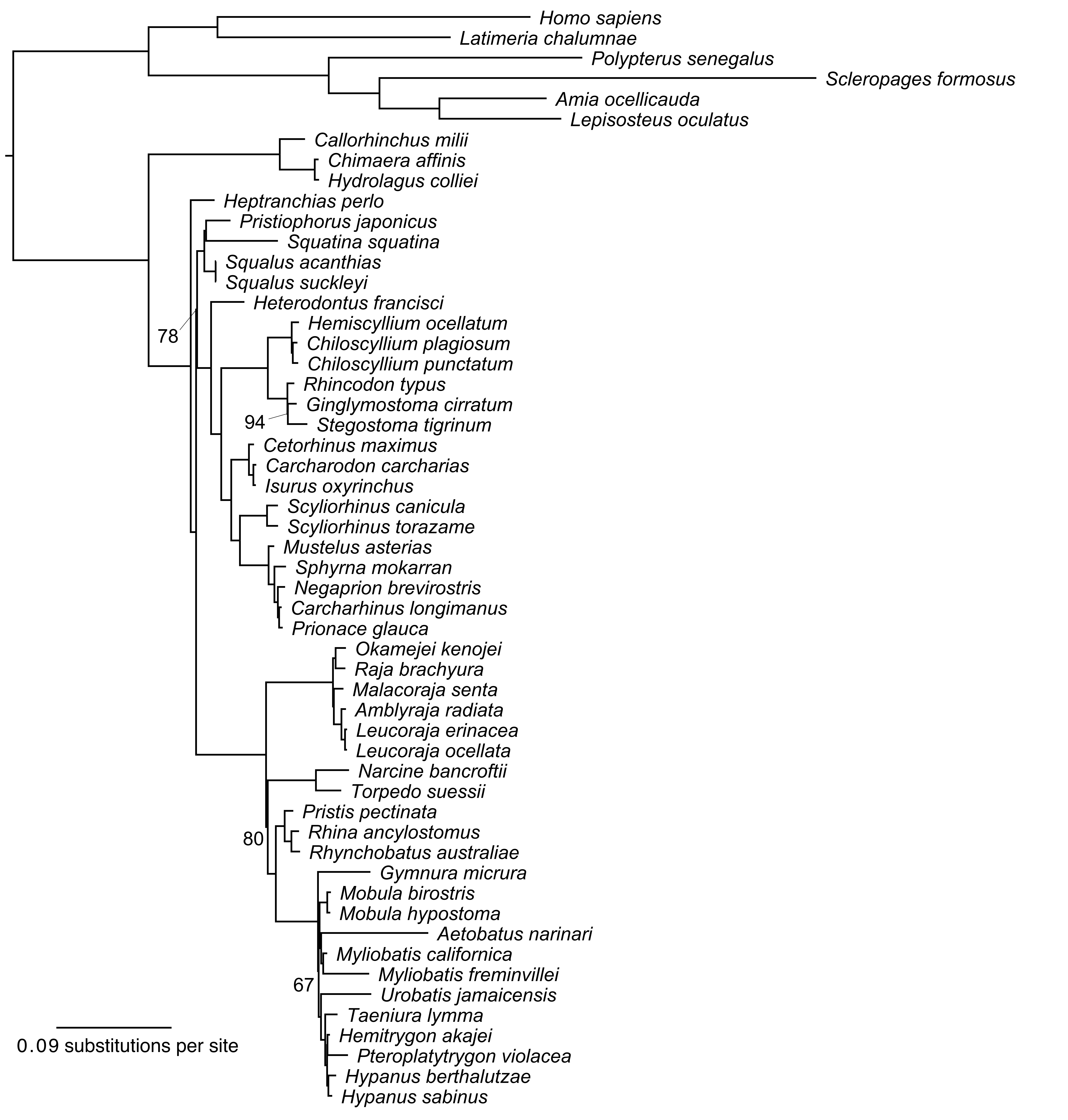

Homo sapiens
Latimeria chalumnae
Polypterus senegalus
Scleropages formosus
Amia ocellicauda
Lepisosteus oculatus
Callorhinchus milii
Chimaera affinis
Hydrolagus colliei
Heptranchias perlo
Pristiophorus japonicus
Squatina squatina
Squalus acanthias
Squalus suckleyi
Heterodontus francisci
78
Hemiscyllium ocellatum
Chiloscyllium plagiosum
Chiloscyllium punctatum
Rhincodon typus
Ginglymostoma cirratum
94
Stegostoma tigrinum
Cetorhinus maximus
Carcharodon carcharias
Isurus oxyrinchus
Scyliorhinus canicula
Scyliorhinus torazame
Mustelus asterias
Sphyrna mokarran
Negaprion brevirostris
Carcharhinus longimanus
Prionace glauca
Okamejei kenojei
Raja brachyura
Malacoraja senta
Amblyraja radiata
Leucoraja erinacea
Leucoraja ocellata
Narcine bancroftii
Torpedo suessii
Pristis pectinata
80
Rhina ancylostomus
Rhynchobatus australiae
Gymnura micrura
Mobula birostris
Mobula hypostoma
Aetobatus narinari
Myliobatis californica
Myliobatis freminvillei
67
Urobatis jamaicensis
Taeniura lymma
Hemitrygon akajei
Pteroplatytrygon violacea
Hypanus berthalutzae
Hypanus sabinus
0.09 substitutions per site

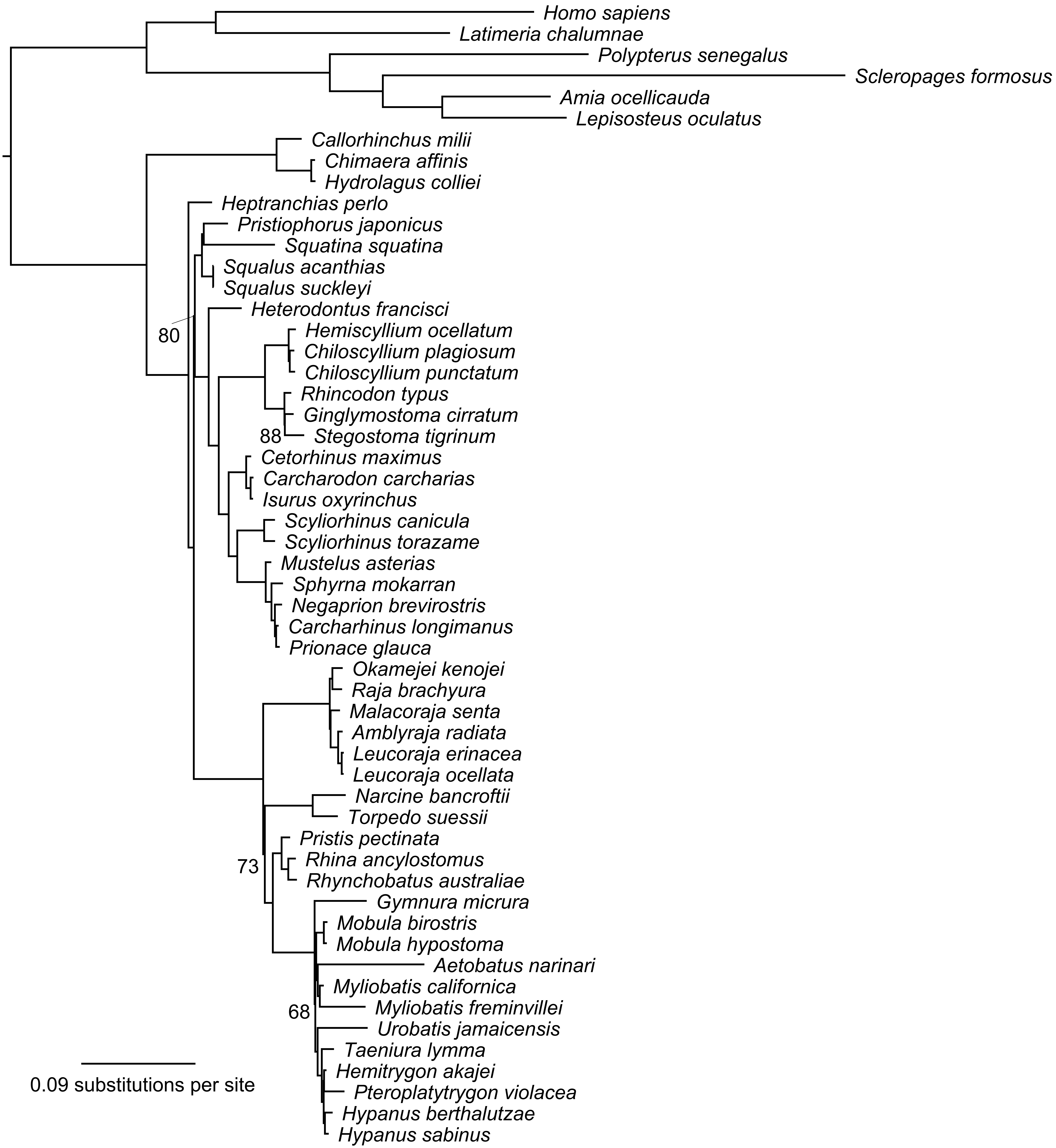
Homo sapiens
Latimeria chalumnae
Polypterus senegalus
Scleropages formosus
Amia ocellicauda
Lepisosteus oculatus
Callorhinchus milii
Chimaera affinis
Hydrolagus colliei
Heptranchias perlo
Pristiophorus japonicus
Squatina squatina
Squalus acanthias
Squalus suckleyi
Heterodontus francisci
80
Hemiscyllium ocellatum
Chiloscyllium plagiosum
Chiloscyllium punctatum
Rhincodon typus
Ginglymostoma cirratum
88
Stegostoma tigrinum
Cetorhinus maximus
Carcharodon carcharias
Isurus oxyrinchus
Scyliorhinus canicula
Scyliorhinus torazame
Mustelus asterias
Sphyrna mokarran
Negaprion brevirostris
Carcharhinus longimanus
Prionace glauca
Okamejei kenojei
Raja brachyura
Malacoraja senta
Amblyraja radiata
Leucoraja erinacea
Leucoraja ocellata
Narcine bancroftii
Torpedo suessii
Pristis pectinata
73
Rhina ancylostomus
Rhynchobatus australiae
Gymnura micrura
Mobula birostris
Mobula hypostoma
Aetobatus narinari
Myliobatis californica
68
Myliobatis freminvillei
Urobatis jamaicensis
Taeniura lymma
Hemitrygon akajei
Pteroplatytrygon violacea
Hypanus berthalutzae
Hypanus sabinus
0.09 substitutions per site

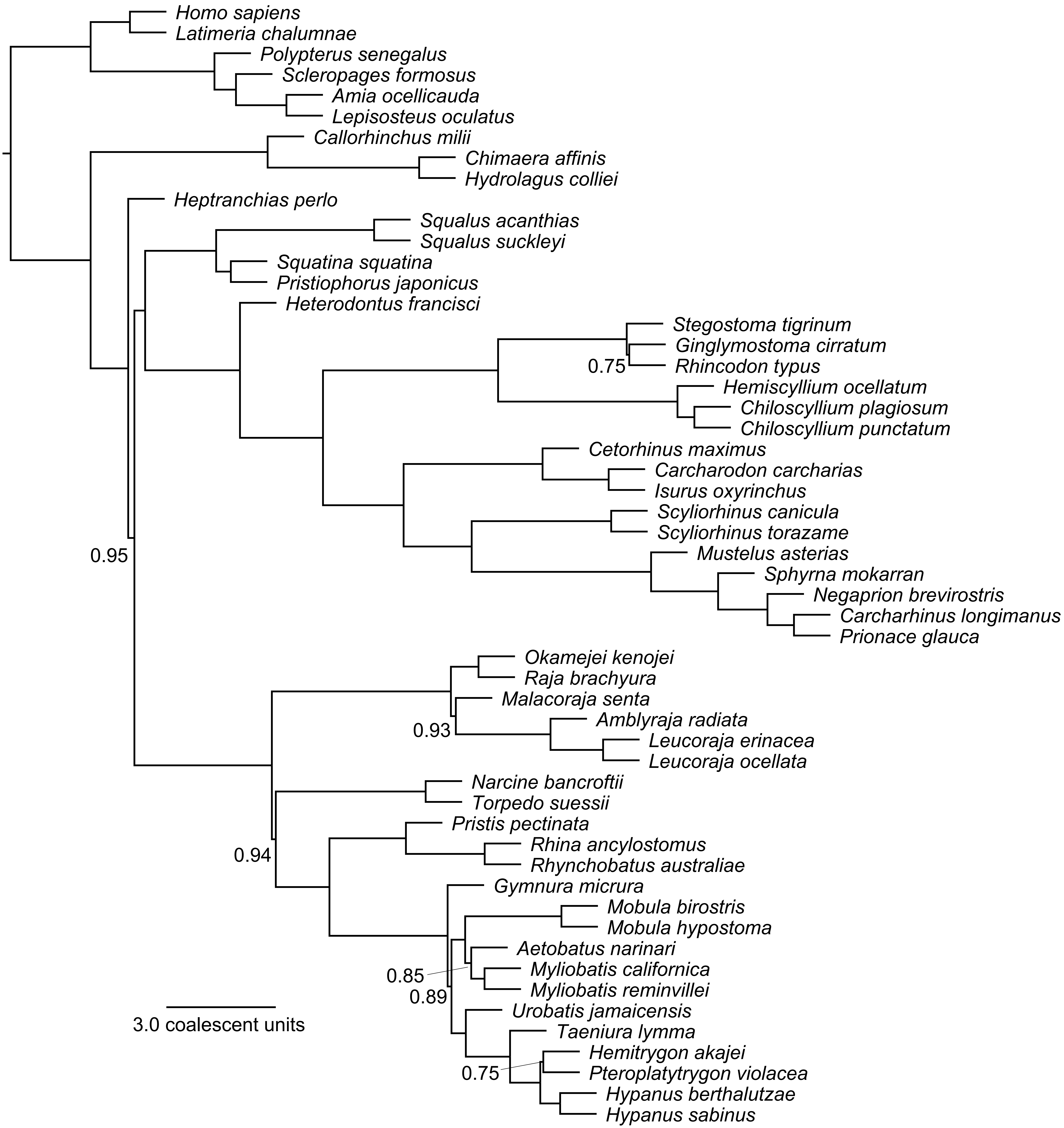
Homo sapiens
Latimeria chalumnae
Polypterus senegalus
Scleropages formosus
Amia ocellicauda
Lepisosteus oculatus
Callorhinchus milii
Chimaera affinis
Hydrolagus colliei
Heptranchias perlo
Squalus acanthias
Squalus suckleyi
Squatina squatina
Pristiophorus japonicus
Heterodontus francisci
Stegostoma tigrinum
Ginglymostoma cirratum
0.75
Rhincodon typus
Hemiscyllium ocellatum
Chiloscyllium plagiosum
Chiloscyllium punctatum
Cetorhinus maximus
Carcharodon carcharias
Isurus oxyrinchus
Scyliorhinus canicula
Scyliorhinus torazame
0.95
Mustelus asterias
Sphyrna mokarran
Negaprion brevirostris
Carcharhinus longimanus
Prionace glauca
Okamejei kenojei
Raja brachyura
Malacoraja senta
Amblyraja radiata
0.93
Leucoraja erinacea
Leucoraja ocellata
Narcine bancroftii
Torpedo suessii
Pristis pectinata
Rhina ancylostomus
0.94
Rhynchobatus australiae
Gymnura micrura
Mobula birostris
Mobula hypostoma
Aetobatus narinari
0.85
Myliobatis californica
0.89
Myliobatis reminvillei
3.0 coalescent units
Urobatis jamaicensis
Taeniura lymma
Hemitrygon akajei
0.75
Pteroplatytrygon violacea
Hypanus berthalutzae
Hypanus sabinus

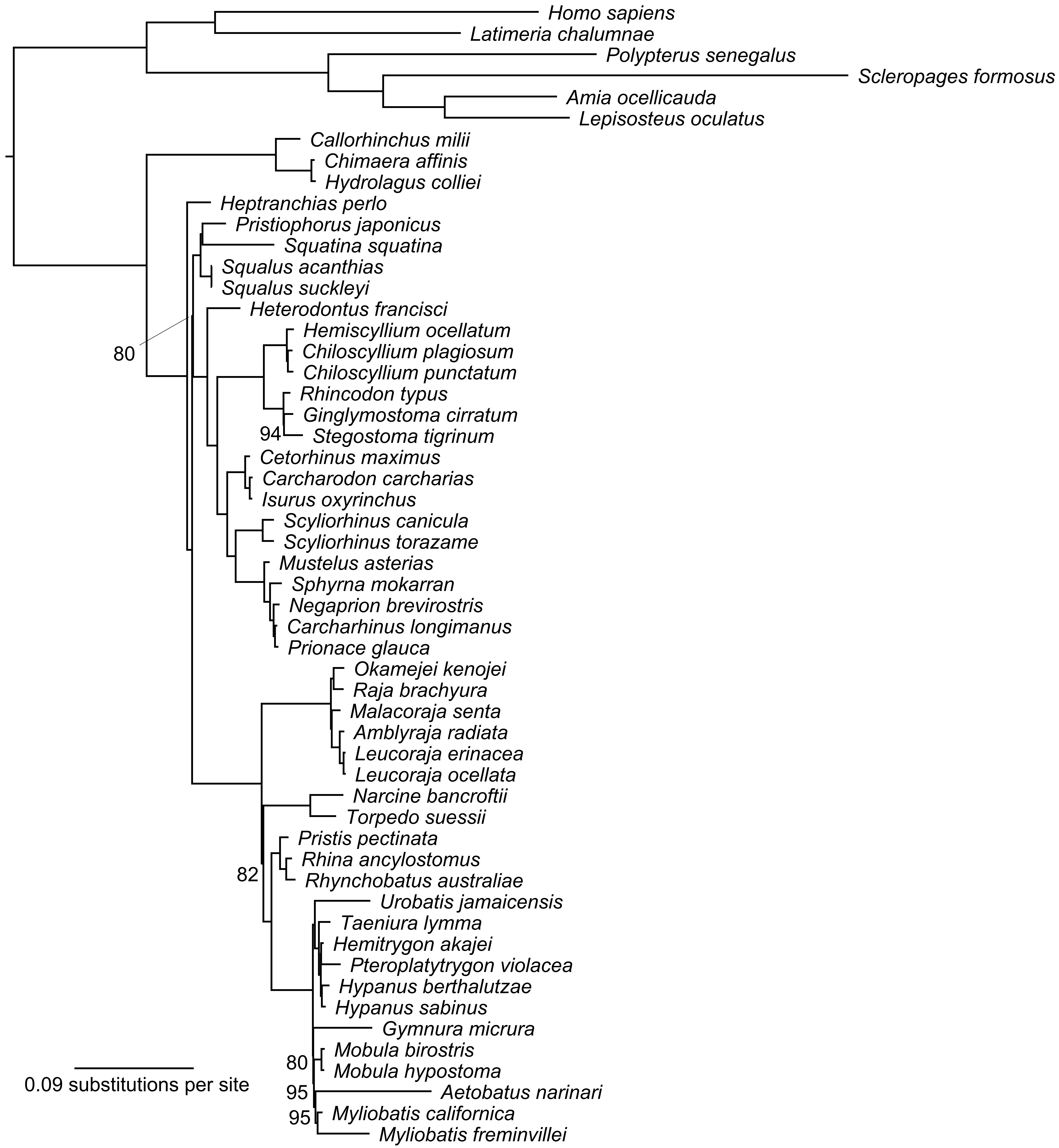
Homo sapiens
Latimeria chalumnae
Polypterus senegalus
Scleropages formosus
Amia ocellicauda
Lepisosteus oculatus
Callorhinchus milii
Chimaera affinis
Hydrolagus colliei
Heptranchias perlo
Pristiophorus japonicus
Squatina squatina
Squalus acanthias
Squalus suckleyi
Heterodontus francisci
80
Hemiscyllium ocellatum
Chiloscyllium plagiosum
Chiloscyllium punctatum
Rhincodon typus
Ginglymostoma cirratum
94
Stegostoma tigrinum
Cetorhinus maximus
Carcharodon carcharias
Isurus oxyrinchus
Scyliorhinus canicula
Scyliorhinus torazame
Mustelus asterias
Sphyrna mokarran
Negaprion brevirostris
Carcharhinus longimanus
Prionace glauca
Okamejei kenojei
Raja brachyura
Malacoraja senta
Amblyraja radiata
Leucoraja erinacea
Leucoraja ocellata
Narcine bancroftii
Torpedo suessii
Pristis pectinata
Rhina ancylostomus
82
Rhynchobatus australiae
Urobatis jamaicensis
Taeniura lymma
Hemitrygon akajei
Pteroplatytrygon violacea
Hypanus berthalutzae
Hypanus sabinus
Gymnura micrura
Mobula birostris
80
Mobula hypostoma
95
Aetobatus narinari
95
Myliobatis californica
Myliobatis freminvillei
0.09 substitutions per site

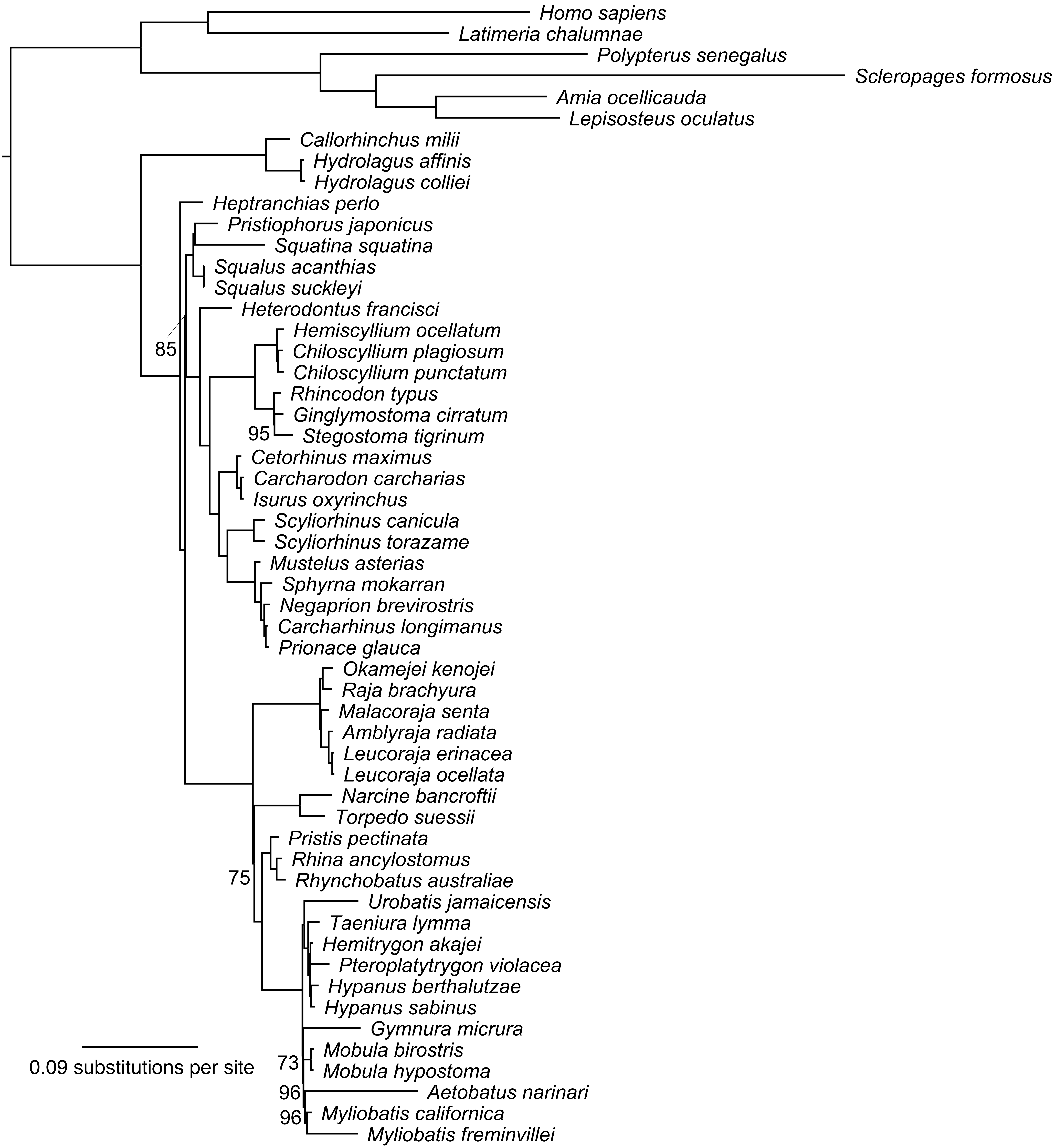
Homo sapiens
Latimeria chalumnae
Polypterus senegalus
Scleropages formosus
Amia ocellicauda
Lepisosteus oculatus
Callorhinchus milii
Hydrolagus affinis
Hydrolagus colliei
Heptranchias perlo
Pristiophorus japonicus
Squatina squatina
Squalus acanthias
Squalus suckleyi
Heterodontus francisci
85
Hemiscyllium ocellatum
Chiloscyllium plagiosum
Chiloscyllium punctatum
Rhincodon typus
Ginglymostoma cirratum
95
Stegostoma tigrinum
Cetorhinus maximus
Carcharodon carcharias
Isurus oxyrinchus
Scyliorhinus canicula
Scyliorhinus torazame
Mustelus asterias
Sphyrna mokarran
Negaprion brevirostris
Carcharhinus longimanus
Prionace glauca
Okamejei kenojei
Raja brachyura
Malacoraja senta
Amblyraja radiata
Leucoraja erinacea
Leucoraja ocellata
Narcine bancroftii
Torpedo suessii
Pristis pectinata
Rhina ancylostomus
75
Rhynchobatus australiae
Urobatis jamaicensis
Taeniura lymma
Hemitrygon akajei
Pteroplatytrygon violacea
Hypanus berthalutzae
Hypanus sabinus
Gymnura micrura
Mobula birostris
73
Mobula hypostoma
96
Aetobatus narinari
96
Myliobatis californica
Myliobatis freminvillei
0.09 substitutions per site

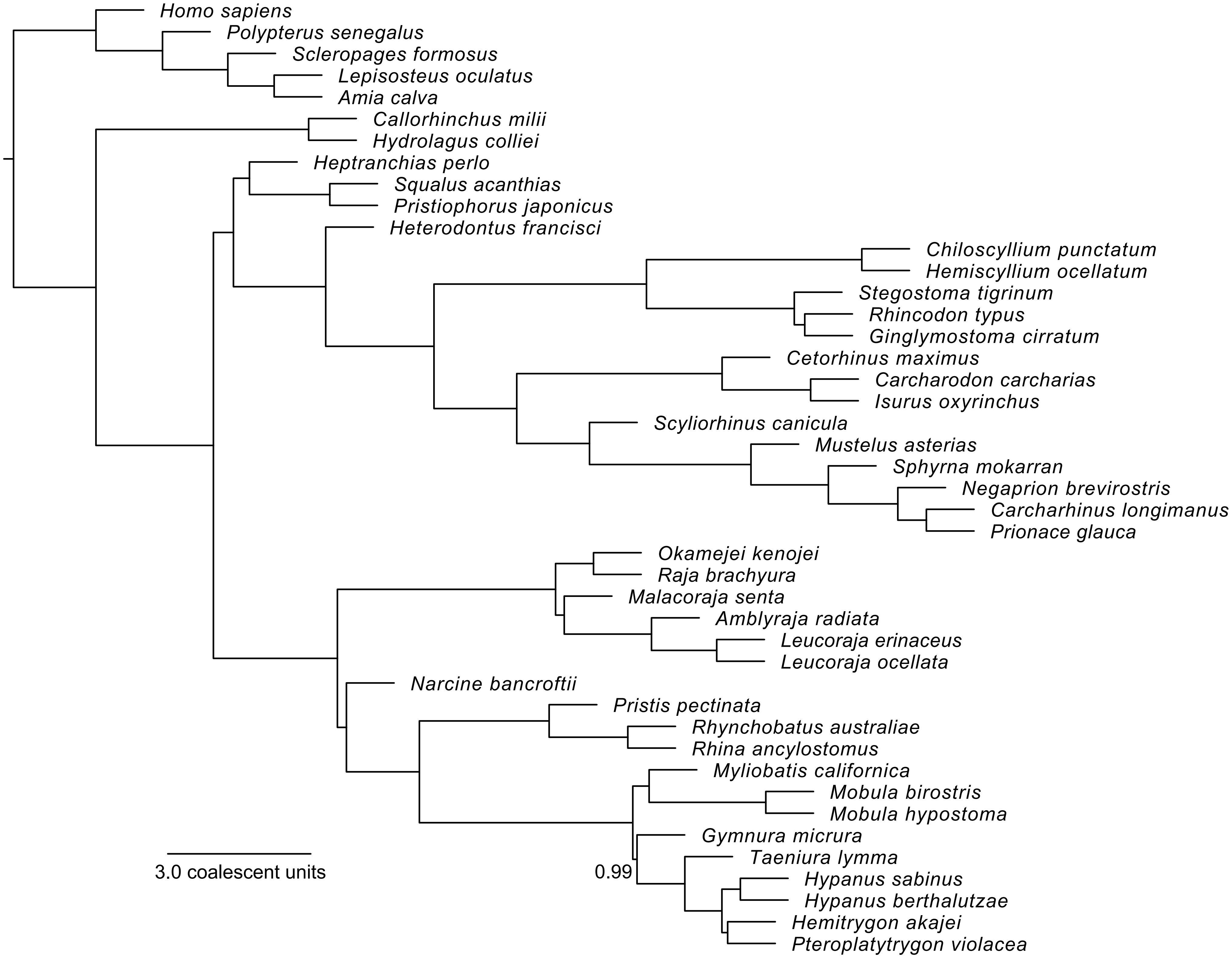
Homo sapiens
Polypterus senegalus
Scleropages formosus
Lepisosteus oculatus
Amia calva
Callorhinchus milii
Hydrolagus colliei
Heptranchias perlo
Squalus acanthias
Pristiophorus japonicus
Heterodontus francisci
Chiloscyllium punctatum
Hemiscyllium ocellatum
Stegostoma tigrinum
Rhincodon typus
Ginglymostoma cirratum
Cetorhinus maximus
Carcharodon carcharias
Isurus oxyrinchus
Scyliorhinus canicula
Mustelus asterias
Sphyrna mokarran
Negaprion brevirostris
Carcharhinus longimanus
Prionace glauca
Okamejei kenojei
Raja brachyura
Malacoraja senta
Amblyraja radiata
Leucoraja erinaceus
Leucoraja ocellata
Narcine bancroftii
Pristis pectinata
Rhynchobatus australiae
Rhina ancylostomus
Myliobatis californica
Mobula birostris
Mobula hypostoma
Gymnura micrura
Taeniura lymma
Hypanus sabinus
Hypanus berthalutzae
Hemitrygon akajei
Pteroplatytrygon violacea
0.99
3.0 coalescent units

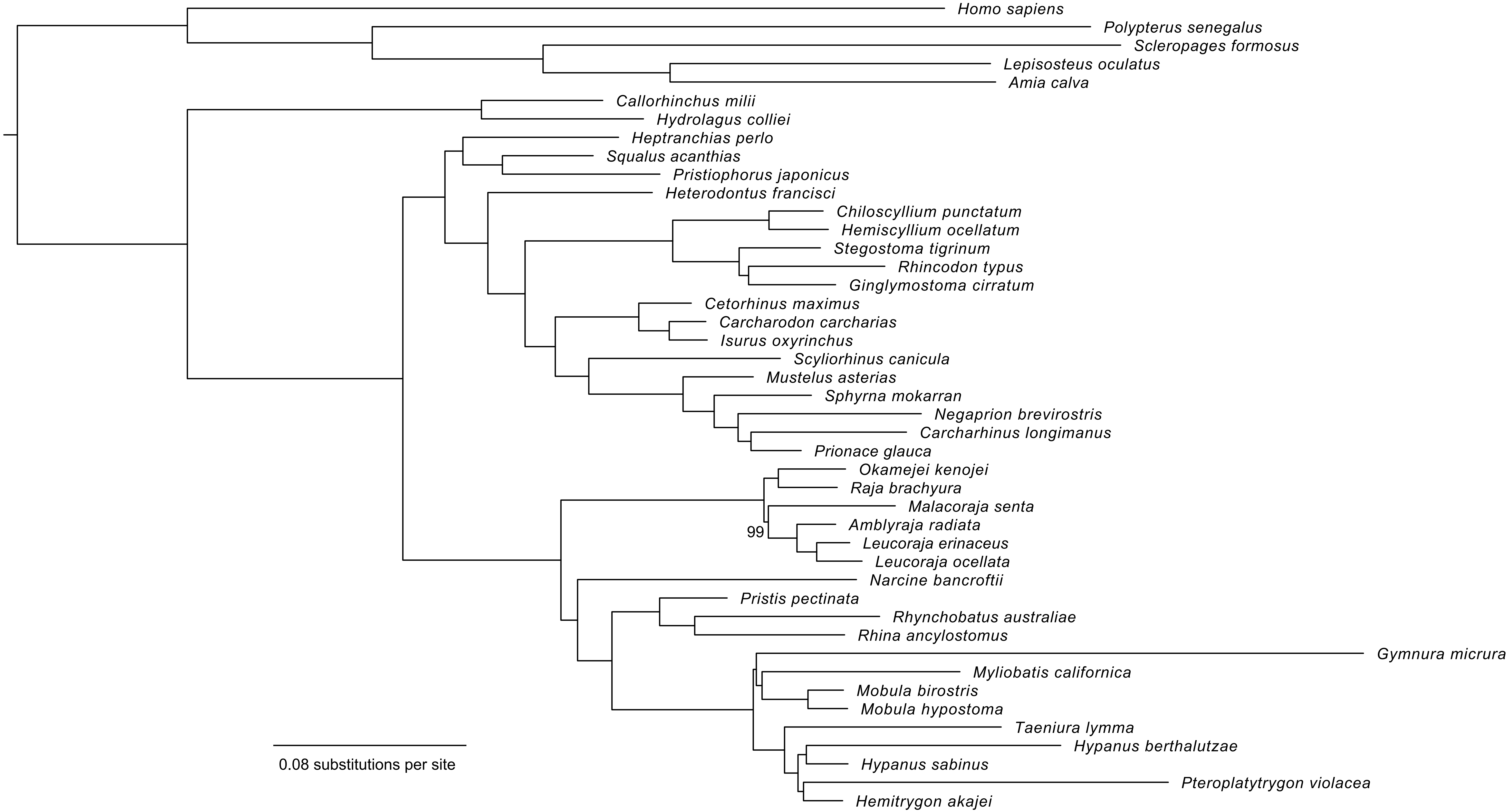
Homo sapiens
Polypterus senegalus
Scleropages formosus
Lepisosteus oculatus
Amia calva
Callorhinchus milii
Hydrolagus colliei
Heptranchias perlo
Squalus acanthias
Pristiophorus japonicus
Heterodontus francisci
Chiloscyllium punctatum
Hemiscyllium ocellatum
Stegostoma tigrinum
Rhincodon typus
Ginglymostoma cirratum
Cetorhinus maximus
Carcharodon carcharias
Isurus oxyrinchus
Scyliorhinus canicula
Mustelus asterias
Sphyrna mokarran
Negaprion brevirostris
Carcharhinus longimanus
Prionace glauca
Okamejei kenojei
Raja brachyura
Malacoraja senta
Amblyraja radiata
99
Leucoraja erinaceus
Leucoraja ocellata
Narcine bancroftii
Pristis pectinata
Rhynchobatus australiae
Rhina ancylostomus
Gymnura micrura
Myliobatis californica
Mobula birostris
Mobula hypostoma
Taeniura lymma
Hypanus berthalutzae
Hypanus sabinus
Pteroplatytrygon violacea
Hemitrygon akajei
0.08 substitutions per site

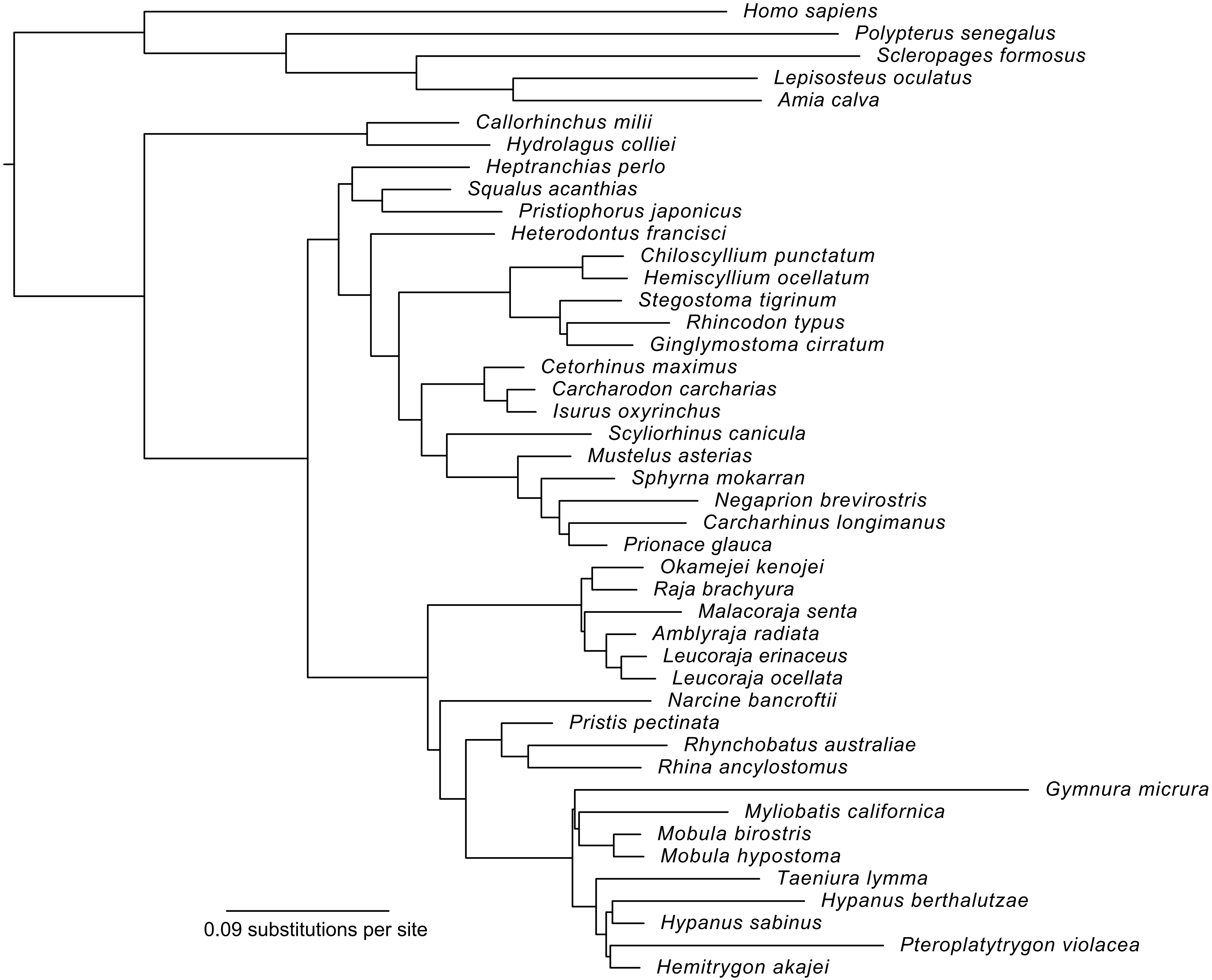
Homo sapiens
Polypterus senegalus
Scleropages formosus
Lepisosteus oculatus
Amia calva
Callorhinchus milii
Hydrolagus colliei
Heptranchias perlo
Squalus acanthias
Pristiophorus japonicus
Heterodontus francisci
Chiloscyllium punctatum
Hemiscyllium ocellatum
Stegostoma tigrinum
Rhincodon typus
Ginglymostoma cirratum
Cetorhinus maximus
Carcharodon carcharias
Isurus oxyrinchus
Scyliorhinus canicula
Mustelus asterias
Sphyrna mokarran
Negaprion brevirostris
Carcharhinus longimanus
Prionace glauca
Okamejei kenojei
Raja brachyura
Malacoraja senta
Amblyraja radiata
Leucoraja erinaceus
Leucoraja ocellata
Narcine bancroftii
Pristis pectinata
Rhynchobatus australiae
Rhina ancylostomus
Gymnura micrura
Myliobatis californica
Mobula birostris
Mobula hypostoma
Taeniura lymma
Hypanus berthalutzae
Hypanus sabinus
Pteroplatytrygon violacea
Hemitrygon akajei
0.09 substitutions per site

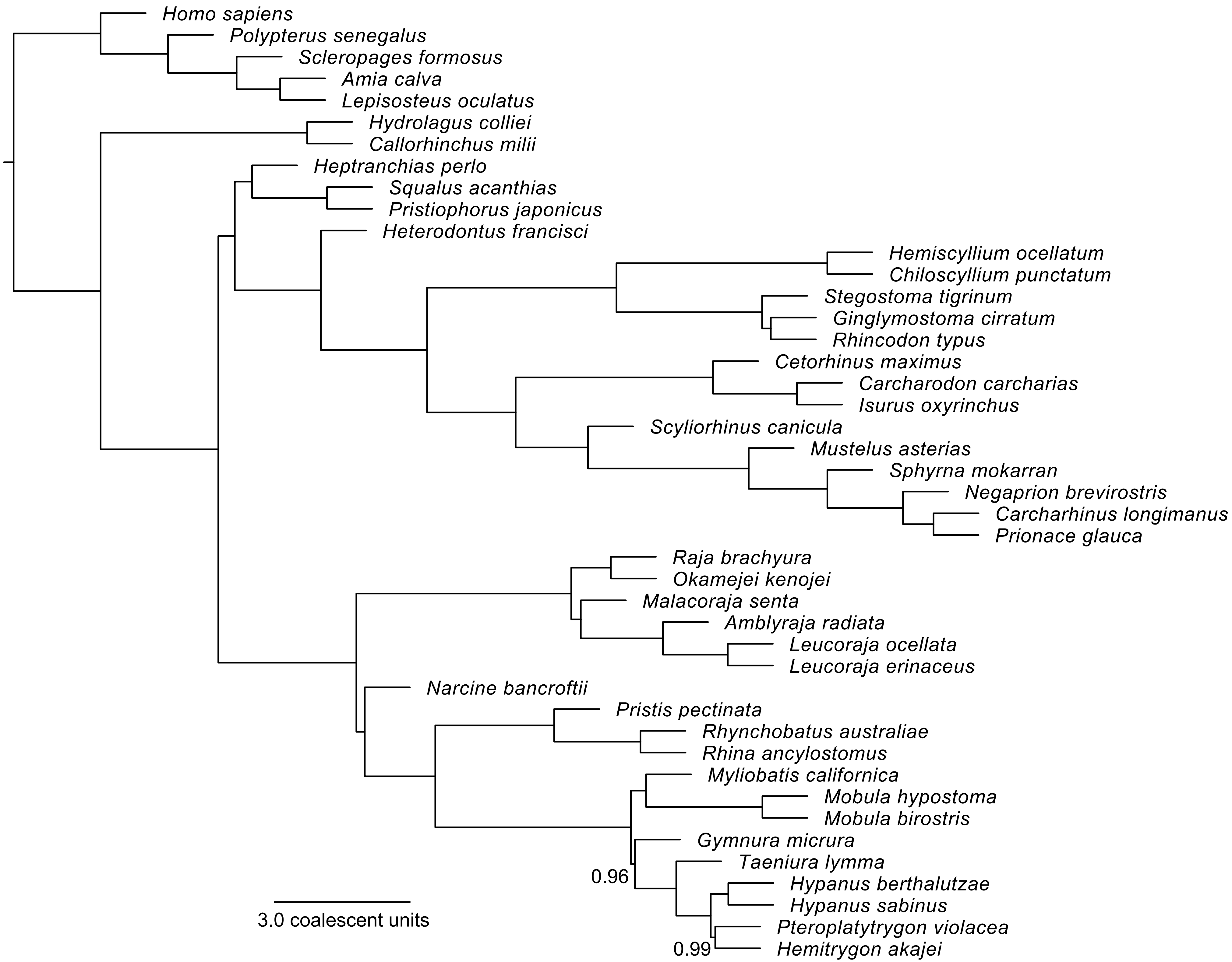

Homo sapiens
Polypterus senegalus
Scleropages formosus
Amia calva
Lepisosteus oculatus
Hydrolagus colliei
Callorhinchus milii
Heptranchias perlo
Squalus acanthias
Pristiophorus japonicus
Heterodontus francisci
Hemiscyllium ocellatum
Chiloscyllium punctatum
Stegostoma tigrinum
Ginglymostoma cirratum
Rhincodon typus
Cetorhinus maximus
Carcharodon carcharias
Isurus oxyrinchus
Scyliorhinus canicula
Mustelus asterias
Sphyrna mokarran
Negaprion brevirostris
Carcharhinus longimanus
Prionace glauca
Raja brachyura
Okamejei kenojei
Malacoraja senta
Amblyraja radiata
Leucoraja ocellata
Leucoraja erinaceus
Narcine bancroftii
Pristis pectinata
Rhynchobatus australiae
Rhina ancylostomus
Myliobatis californica
Mobula hypostoma
Mobula birostris
Gymnura micrura
Taeniura lymma
Hypanus berthalutzae
Hypanus sabinus
Pteroplatytrygon violacea
Hemitrygon akajei
0.96
0.99
3.0 coalescent units

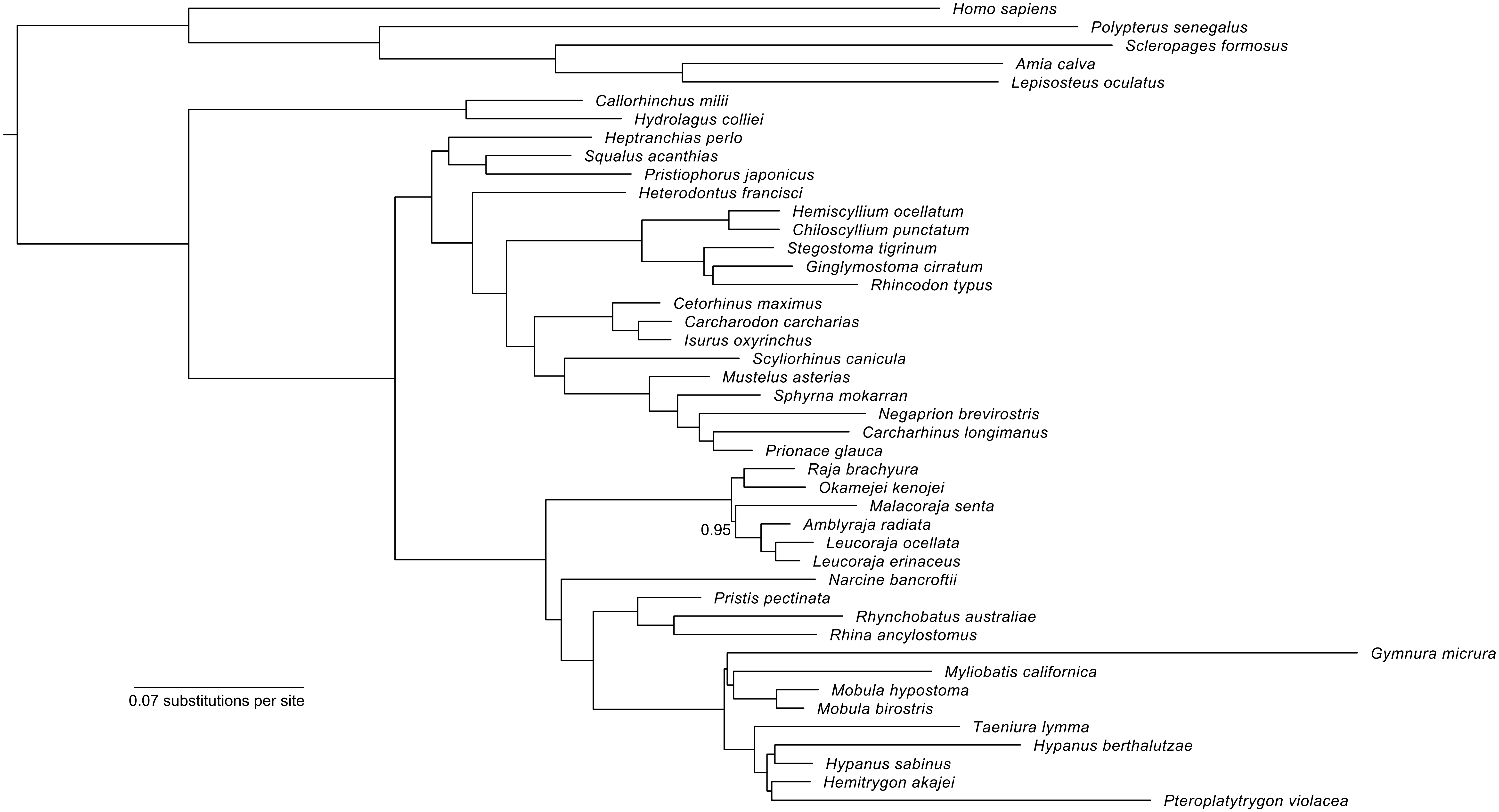
Homo sapiens
Polypterus senegalus
Scleropages formosus
Amia calva
Lepisosteus oculatus
Callorhinchus milii
Hydrolagus colliei
Heptranchias perlo
Squalus acanthias
Pristiophorus japonicus
Heterodontus francisci
Hemiscyllium ocellatum
Chiloscyllium punctatum
Stegostoma tigrinum
Ginglymostoma cirratum
Rhincodon typus
Cetorhinus maximus
Carcharodon carcharias
Isurus oxyrinchus
Scyliorhinus canicula
Mustelus asterias
Sphyrna mokarran
Negaprion brevirostris
Carcharhinus longimanus
Prionace glauca
Raja brachyura
Okamejei kenojei
Malacoraja senta
0.95
Amblyraja radiata
Leucoraja ocellata
Leucoraja erinaceus
Narcine bancroftii
Pristis pectinata
Rhynchobatus australiae
Rhina ancylostomus
Gymnura micrura
Myliobatis californica
Mobula hypostoma
Mobula birostris
Taeniura lymma
Hypanus berthalutzae
Hypanus sabinus
Hemitrygon akajei
Pteroplatytrygon violacea
0.07 substitutions per site

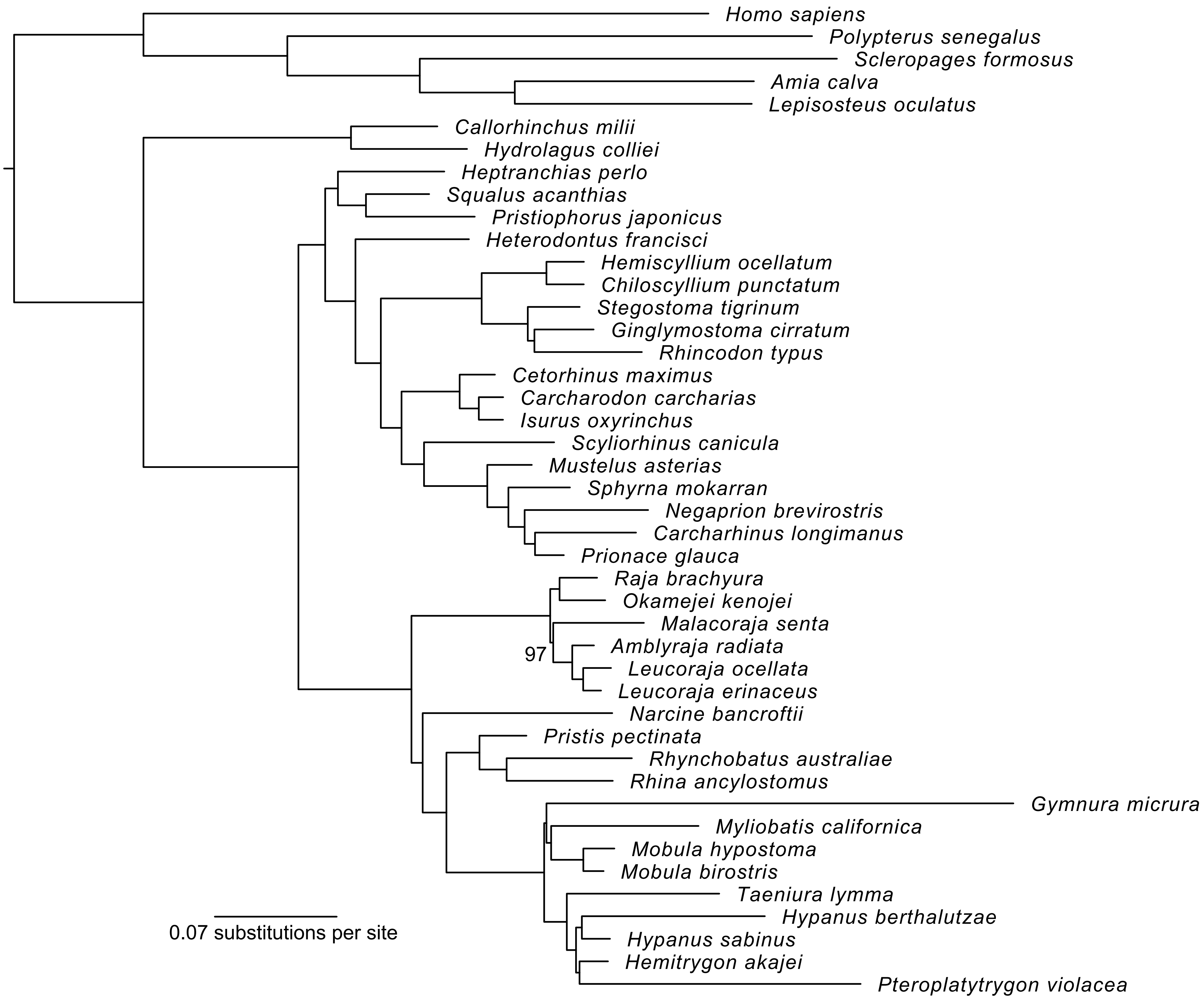

Homo sapiens
Polypterus senegalus
Scleropages formosus
Amia calva
Lepisosteus oculatus
Callorhinchus milii
Hydrolagus colliei
Heptranchias perlo
Squalus acanthias
Pristiophorus japonicus
Heterodontus francisci
Hemiscyllium ocellatum
Chiloscyllium punctatum
Stegostoma tigrinum
Ginglymostoma cirratum
Rhincodon typus
Cetorhinus maximus
Carcharodon carcharias
Isurus oxyrinchus
Scyliorhinus canicula
Mustelus asterias
Sphyrna mokarran
Negaprion brevirostris
Carcharhinus longimanus
Prionace glauca
Raja brachyura
Okamejei kenojei
Malacoraja senta
97
Amblyraja radiata
Leucoraja ocellata
Leucoraja erinaceus
Narcine bancroftii
Pristis pectinata
Rhynchobatus australiae
Rhina ancylostomus
Gymnura micrura
Myliobatis californica
Mobula hypostoma
Mobula birostris
Taeniura lymma
Hypanus berthalutzae
Hypanus sabinus
Hemitrygon akajei
Pteroplatytrygon violacea
0.07 substitutions per site

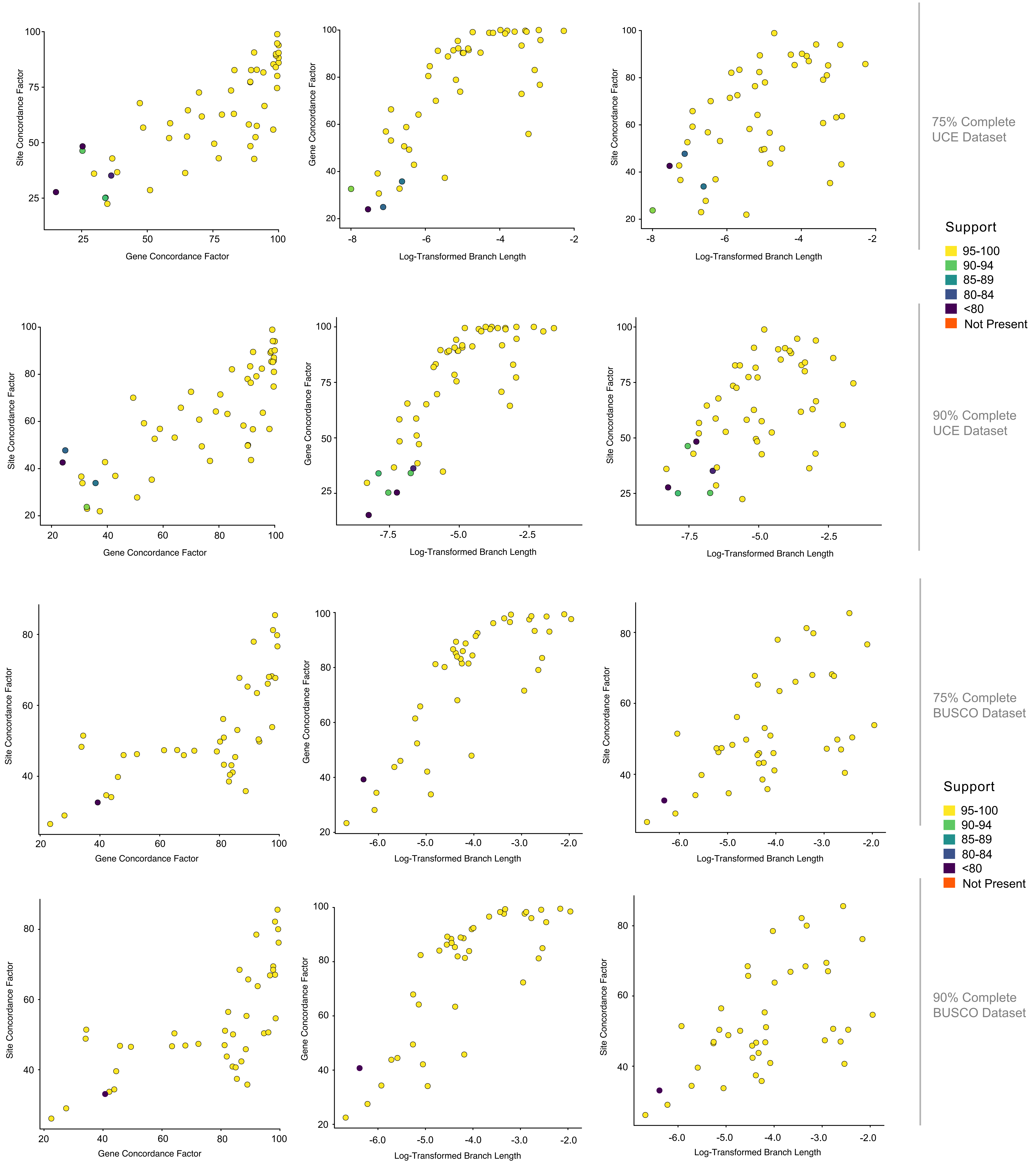
Site Concordance Factor
Gene Concordance Factor
Log-Transformed Branch Length
75% Complete
UCE Dataset
90% Complete
UCE Dataset
75% Complete
BUSCO Dataset
90% Complete
BUSCO Dataset
Support
95-100
90-94
85-89
80-84
<80
Not Present

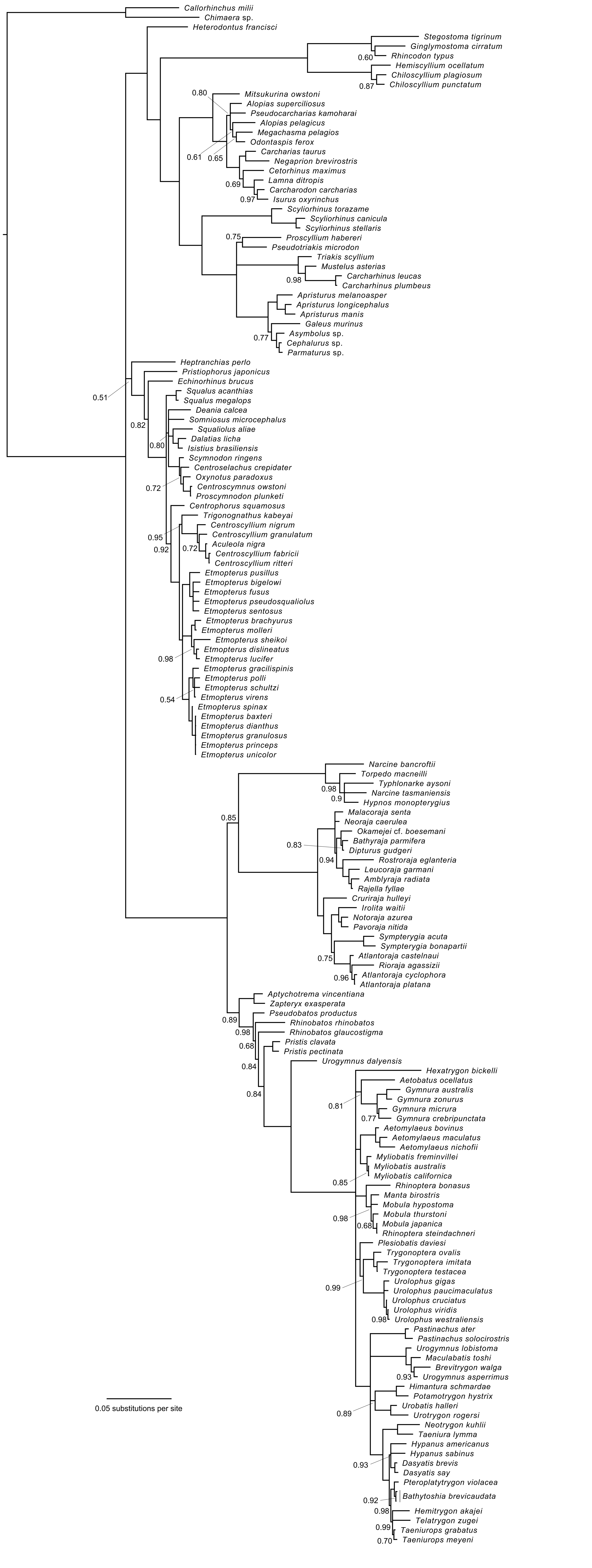
Callorhinchus milii
Chimaera sp.
Heterodontus francisci
Stegostoma tigrinum
Ginglymostoma cirratum
Rhincodon typus
Hemiscyllium ocellatum
Chiloscyllium plagiosum
Chiloscyllium punctatum
Mitsukurina owstoni
Alopias superciliosus
Pseudocarcharias kamoharai
Alopias pelagicus
Megachasma pelagios
Odontaspis ferox
Carcharias taurus
Negaprion brevirostris
Cetorhinus maximus
Lamna ditropis
Carcharodon carcharias
Isurus oxyrinchus
Scyliorhinus torazame
Scyliorhinus canicula
Scyliorhinus stellaris
Proscyllium habereri
Pseudotriakis microdon
Triakis scyllium
Mustelus asterias
Carcharhinus leucas
Carcharhinus plumbeus
Apristurus melanoasper
Apristurus longicephalus
Apristurus manis
Galeus murinus
Asymbolus sp.
Cephalurus sp.
Parmaturus sp.
Heptranchias perlo
Pristiophorus japonicus
Echinorhinus brucus
Squalus acanthias
Squalus megalops
Deania calcea
Somniosus microcephalus
Squaliolus aliae
Dalatias licha
Isistius brasiliensis
Scymnodon ringens
Centroselachus crepidater
Oxynotus paradoxus
Centroscymnus owstoni
Proscymnodon plunketi
Centrophorus squamosus
Trigonognathus kabeyai
Centroscyllium nigrum
Centroscyllium granulatum
Aculeola nigra
Centroscyllium fabricii
Centroscyllium ritteri
Etmopterus pusillus
Etmopterus bigelowi
Etmopterus fusus
Etmopterus pseudosqualiolus
Etmopterus sentosus
Etmopterus brachyurus
Etmopterus molleri
Etmopterus sheikoi
Etmopterus dislineatus
Etmopterus lucifer
Etmopterus gracilispinis
Etmopterus polli
Etmopterus schultzi
Etmopterus virens
Etmopterus spinax
Etmopterus baxteri
Etmopterus dianthus
Etmopterus granulosus
Etmopterus princeps
Etmopterus unicolor
Narcine bancroftii
Torpedo macneilli
Typhlonarke aysoni
Narcine tasmaniensis
Hypnos monopterygius
Malacoraja senta
Neoraja caerulea
Okamejei cf. boesemani
Bathyraja parmifera
Dipturus gudgeri
Rostroraja eglanteria
Leucoraja garmani
Amblyraja radiata
Rajella fyllae
Cruriraja hulleyi
Irolita waitii
Notoraja azurea
Pavoraja nitida
Sympterygia acuta
Sympterygia bonapartii
Atlantoraja castelnaui
Rioraja agassizii
Atlantoraja cyclophora
Atlantoraja platana
Aptychotrema vincentiana
Zapteryx exasperata
Pseudobatos productus
Rhinobatos rhinobatos
Rhinobatos glaucostigma
Pristis clavata
Pristis pectinata
Urogymnus dalyensis
Hexatrygon bickelli
Aetobatus ocellatus
Gymnura australis
Gymnura zonurus
Gymnura micrura
Gymnura crebripunctata
Aetomylaeus bovinus
Aetomylaeus maculatus
Aetomylaeus nichofii
Myliobatis freminvillei
Myliobatis australis
Myliobatis californica
Rhinoptera bonasus
Manta birostris
Mobula hypostoma
Mobula thurstoni
Mobula japanica
Rhinoptera steindachneri
Plesiobatis daviesi
Trygonoptera ovalis
Trygonoptera imitata
Trygonoptera testacea
Urolophus gigas
Urolophus paucimaculatus
Urolophus cruciatus
Urolophus viridis
Urolophus westraliensis
Pastinachus ater
Pastinachus solocirostris
Urogymnus lobistoma
Maculabatis toshi
Brevitrygon walga
Urogymnus asperrimus
Himantura schmardae
Potamotrygon hystrix
Urobatis halleri
Urotrygon rogersi
Neotrygon kuhlii
Taeniura lymma
Hypanus americanus
Hypanus sabinus
Dasyatis brevis
Dasyatis say
Pteroplatytrygon violacea
Bathytoshia brevicaudata
Hemitrygon akajei
Telatrygon zugei
Taeniurops grabatus
Taeniurops meyeni
0.05 substitutions per site

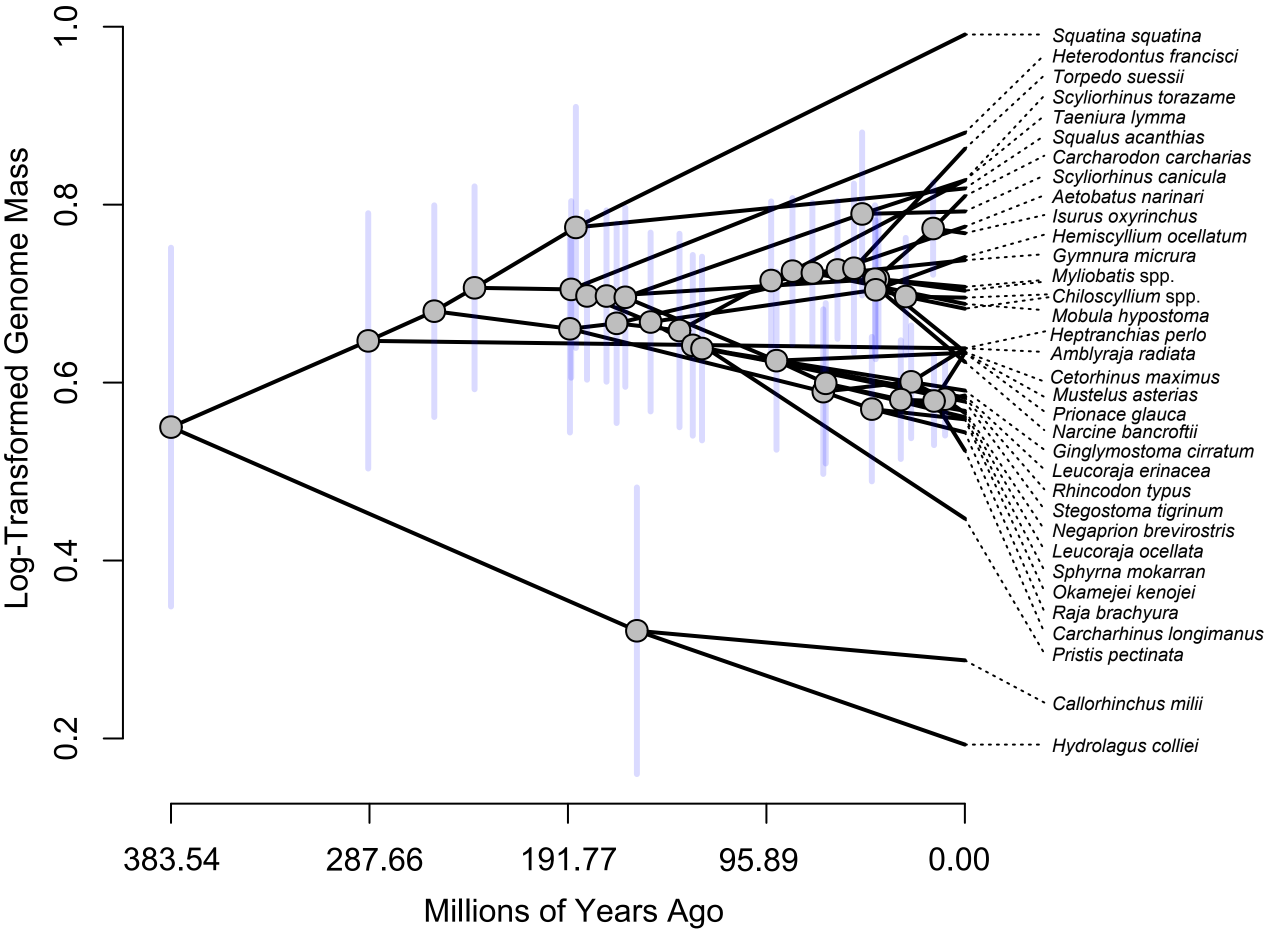

Log-Transformed Genome Mass
1.0
0.8
0.6
0.4
0.2
383.54
287.66
191.77
95.89
0.00
Millions of Years Ago
Squatina squatina
Heterodontus francisci
Torpedo suessii
Scyliorhinus torazame
Taeniura lymma
Squalus acanthias
Carcharodon carcharias
Scyliorhinus canicula
Aetobatus narinari
Isurus oxyrinchus
Hemiscyllium ocellatum
Gymnura micrura
Myliobatis spp.
Chiloscyllium spp.
Mobula hypostoma
Heptranchias perlo
Amblyraja radiata
Cetorhinus maximus
Mustelus asterias
Prionace glauca
Narcine bancroftii
Ginglymostoma cirratum
Leucoraja erinacea
Rhincodon typus
Stegostoma tigrinum
Negaprion brevirostris
Leucoraja ocellata
Sphyrna mokarran
Okamejei kenojei
Raja brachyura
Carcharhinus longimanus
Pristis pectinata
Callorhinchus milii
Hydrolagus colliei
